# Screening for genes that accelerate the epigenetic ageing clock in humans reveals a role for the H3K36 methyltransferase NSD1

**DOI:** 10.1101/545830

**Authors:** Daniel E. Martin-Herranz, Erfan Aref-Eshghi, Marc Jan Bonder, Thomas M. Stubbs, Oliver Stegle, Bekim Sadikovic, Wolf Reik, Janet M. Thornton

## Abstract

**Background:** Epigenetic clocks are mathematical models that predict the biological age of an individual using DNA methylation data, and which have emerged in the last few years as the most accurate biomarkers of the ageing process. However, little is known about the molecular mechanisms that control the rate of such clocks. Here, we have examined the human epigenetic clock in patients with a variety of developmental disorders, harbouring mutations in proteins of the epigenetic machinery.

**Results:** Using the Horvath epigenetic clock, we performed an unbiased screen for epigenetic age acceleration (EAA) in the blood of these patients. We demonstrate that loss-of-function mutations in the H3K36 histone methyltransferase NSD1, which cause Sotos syndrome, substantially accelerate epigenetic ageing. Furthermore, we show that the normal ageing process and Sotos syndrome share methylation changes and the genomic context in which they occur. Finally, we found that the Horvath clock CpG sites are characterised by a higher Shannon methylation entropy when compared with the rest of the genome, which is dramatically decreased in Sotos syndrome patients.

**Conclusions:** These results suggest that the H3K36 methylation machinery is a key component of the *epigenetic maintenance system* in humans, which controls the rate of epigenetic ageing, and this role seems to be conserved in model organisms. Our observations provide novel insights into the mechanisms behind the epigenetic ageing clock and we expect will shed light on the different processes that erode the human epigenetic landscape during ageing.

## BACKGROUND

Ageing is normally defined as the time-dependent functional decline which increases vulnerability to common diseases and death in most organisms [1]. However, the molecular processes that drive the emergence of age-related diseases are only beginning to be elucidated. With the passage of time, dramatic and complex changes accumulate in the epigenome of cells, from yeast to humans, pinpointing epigenetic alterations as one of the hallmarks of ageing [1–4].

Our understanding of the ageing process has historically been hampered by the lack of tools to accurately measure it. In recent years, epigenetic clocks have emerged as powerful biomarkers of the ageing process across mammals [5, 6], including humans [7–9], mouse [10–14], dogs and wolves [15] and humpback whales [16]. Epigenetic clocks are mathematical models that are trained to predict chronological age using the DNA methylation status of a small number of CpG sites in the genome. The most widely used multi-tissue epigenetic clock in humans was developed by Steve Horvath in 2013 [8]. Interestingly, deviations of the epigenetic (biological) age from the expected chronological age (aka epigenetic age acceleration or EAA) have been associated with many conditions in humans, including time-to-death [17, 18], HIV infection [19], Down syndrome [20], obesity [21], Werner syndrome[22] and Huntington’s disease [23]. In mice, the epigenetic clock is slowed down by dwarfism and calorie restriction [11–14, 24] and is accelerated by ovariectomy and high fat diet [10, 13]. Furthermore, *in vitro* reprogramming of somatic cells into iPSCs reduces epigenetic age to values close to zero both in humans [8] and mice [11, 14], which opens the door to potential rejuvenation therapies [25, 26]

Epigenetic clocks can be understood as a proxy to quantify the changes of the epigenome with age. However, little is known about the molecular mechanisms that determine the rate of these clocks. Steve Horvath proposed that the multi-tissue epigenetic clock captures the workings of an *epigenetic maintenance system* [8]. Recent GWAS studies have found several genetic variants associated with epigenetic age acceleration in genes such as *TERT* (the catalytic subunit of telomerase) [27], *DHX57* (an ATP-dependent RNA helicase) [28] or *MLST8* (a subunit of both mTORC1 and mTORC2 complexes) [28]. Nevertheless, to our knowledge no genetic variants in epigenetic modifiers have been found and the molecular nature of this hypothetical system is unknown to this date.

We decided to take a reverse genetics approach and look at the behaviour of the epigenetic clock in patients with developmental disorders, many of which harbour mutations in proteins of the epigenetic machinery [29, 30]. We performed an unbiased screen for epigenetic age acceleration and found that Sotos syndrome accelerates epigenetic ageing, potentially revealing a role of H3K36 methylation maintenance in the regulation of the rate of the epigenetic clock.

## RESULTS

### Screening for epigenetic age acceleration (EAA) is improved when correcting for batch effects

The main goal of this study is to identify genes, mainly components of the epigenetic machinery, that can affect the rate of epigenetic ageing in humans (as measured by Horvath’s epigenetic clock) [8]. For this purpose, we conducted an unbiased screen for epigenetic age acceleration (EAA) in samples from patients with developmental disorders that we could access and for which genome-wide DNA methylation data was available (Table 1, Additional file 2). All the DNA methylation data were generated from blood using the Illumina HumanMethylation450 array (450K array).

**Table 1.** Overview of the developmental disorders that were included in the screening (total N = 367) after quality control (QC) and filtering (see Methods and Fig. 1a).

| Developmental disorder | Gene(s) involved | Gene(s) function | Molecular cause | N | Age range (years) |
| --- | --- | --- | --- | --- | --- |
| Angelman | *UBE3A* | Ubiquitin protein ligase E3A | Imprinting, mutation | 14 | 1 to 55 |
| Autism spectrum disorder (ASD) | - | - | - | 119 | 1.83 to 35.16 |
| Alpha thalassemia/mental retardation X-linked syndrome (ATR-X) | *ATRX* | Chromatin remodelling | Mutation | 15 | 0.7 to 27 |
| Claes-Jensen | *KDM5C* | H3K4 demethylase | Mutation | 10 | 2 to 42 |
| Coffin-Lowry | *RPS6KA3* | Serine/threonine kinase | Mutation | 10 | 1.3 to 22.8 |
| Floating-Harbour | *SRCAP* | Chromatin remodelling | Mutation | 17 | 4 to 42 |
| Fragile X syndrome (FXS) | *FMR1* | Translational control | Mutation (CGG expansion) | 32 | 0.08 to 48 |
| Kabuki | *KMT2D* | H3K4 methyltransferase | Mutation | 46 | 0 to 24.1 |
| Noonan | *PTPN11, RAF1, SOS1* | RAS/MAPK signalling | Mutation | 15,11,14 | 0.2 to 49 |
| Rett | *MECP2* | Transcriptional repression | Mutation | 15 | 1 to 34 |
| Saethre-Chotzen | *TWIST1* | Transcription factor | Mutation | 22 | 0 to 38 |
| Sotos | *NSD1* | H3K36 methyltransferase | Mutation | 20 | 1.6 to 41 |
| Weaver | *EZH2* | H3K27 methyltransferase | Mutation | 7 | 2.58 to 43 |

The main step in the screening methodology is to compare the EAA distribution for the samples with a given developmental disorder against a robust control (Fig. 1a). In our case, the control set was obtained from human blood samples in a healthy population of individuals that matched the age range of the developmental disorder samples (Additional file 3). Given that the EAA reflects deviations between the epigenetic (biological) age and the chronological age of a sample, we would expect the EAA distributions of the controls to be centred around zero, which is equivalent to the situation when the median absolute error (MAE) of the model prediction is close to zero (see Methods). This was not the case for the samples obtained from several control batches (Additional file 1: Figure S1A, Additional file 1: Figure S1B), both in the case of EAA models with and without cell composition correction (CCC). It is worth noting that these results were obtained even after applying the internal normalisation step against a blood gold-standard suggested by Horvath [8]. Therefore, we hypothesised that part of the deviations observed might be caused by technical variance that was affecting epigenetic age predictions in the different batches.

**Fig. 1.**
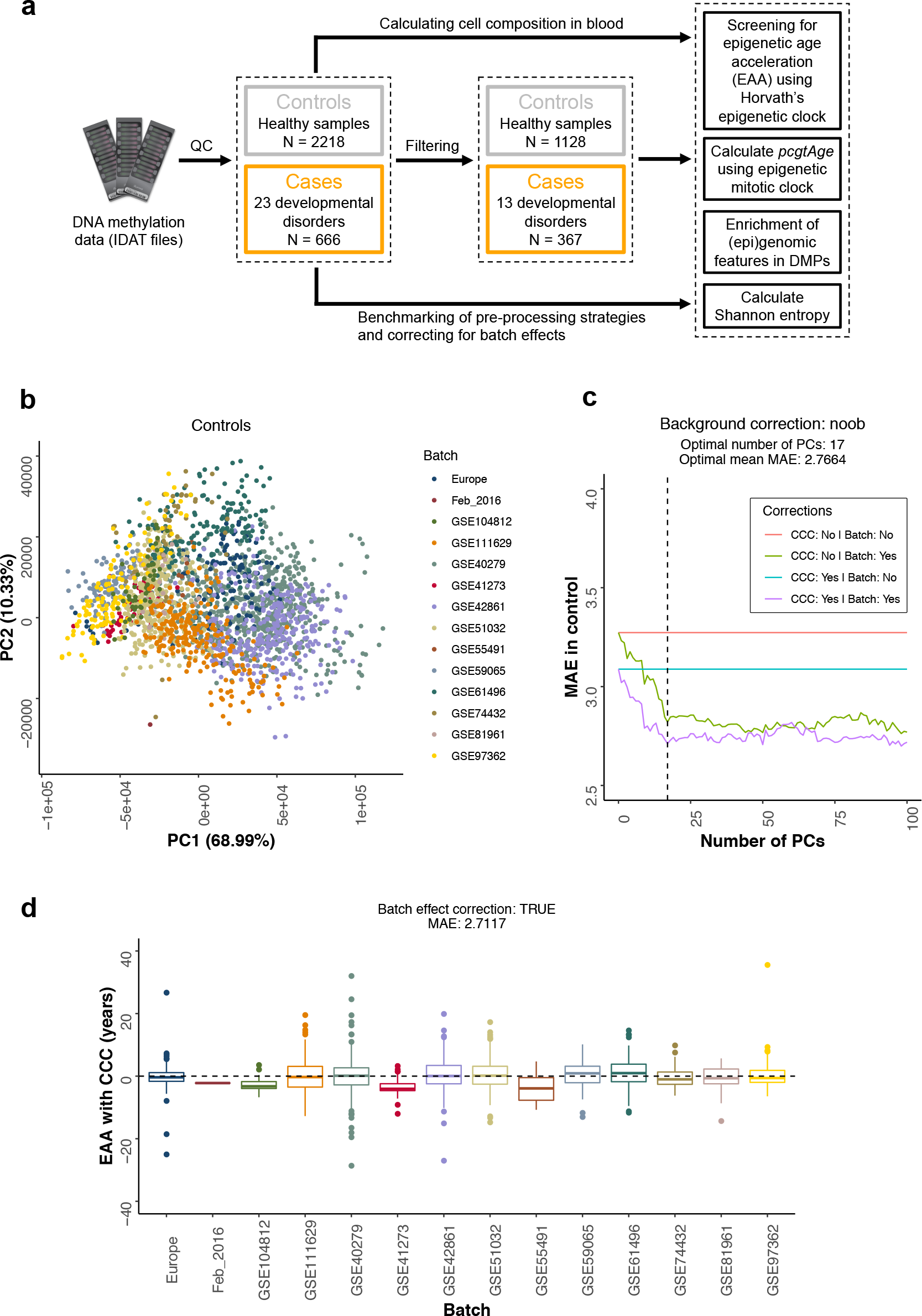
Screening for epigenetic age acceleration (EAA) is improved when correcting for batch effects. **a.** Flow diagram that portrays an overview of the different analyses that are carried out in the raw DNA methylation data (IDAT files) from human blood for cases (developmental disorders samples) and controls (healthy samples). The control samples are filtered to match the age range of the cases (0-55 years). The cases are filtered based on the number of ‘adult’ samples available (for each disorder, at least 5 samples, with 2 of them with an age ≥ 20 years). More details can be found in the Methods. QC: quality control. DMPs: differentially methylated positions. **b.** Scatterplot showing the values of the first two principal components (PCs) for the control samples after performing PCA on the control probes of the 450K arrays. Each point corresponds to a different control sample and the colours represent the different batches. The different batches cluster together in the PCA space, showing that the control probes indeed capture technical variation. Please note that all the PCA calculations were done with more samples from cases and controls than those that were included in the final screening since it was performed before the filtering step (see Methods and Fig. 1a). **c.** Plot showing how the median absolute error (MAE) of the prediction in the control samples, that should tend to zero, is reduced when the PCs capturing the technical variation are included as part of the modelling strategy (see Methods). The dashed line represents the optimal number of PCs (17) that was finally used. The optimal mean MAE is calculated as the average MAE between the green and purple lines. CCC: cell composition correction. **d.** Distribution of the EAA with cell composition correction (CCC) for the different control batches, after applying batch effect correction.

We decided to correct for the potential batch effects by making use of the control probes present on the 450K array, which have been shown to carry information about unwanted variation from a technical source (i.e. technical variance) [31–33]. Performing principal components analysis (PCA) on the raw intensities of the control probes showed that the first two components (PCs) capture the batch structure in both controls (Fig. 1b) and cases (Additional file 1: Figure S1C). Including the first 17 PCs as part of the EAA modelling strategy (see Methods), which together accounted for 98.06% of the technical variance in controls and cases (Additional file 1: Figure S1D), significantly reduced the median absolute error (MAE) of the predictions in the controls (MAE without CCC = 2.8211 years, MAE with CCC = 2.7117 years, mean MAE = 2.7664 years, Fig. 1c). These values are below the original MAE reported by Horvath in his test set (3.6 years) [8].

Finally, deviations from a median EAA close to zero in some of the control batches after batch effect correction (Fig. 1d, Additional file 1: Figure S1E) could be explained by other variables, such as a small batch size or an overrepresentation of young samples (Additional file 1: Figure S1F). The latter is a consequence of the fact that Horvath’s model underestimates the epigenetic ages of older samples, a phenomenon which has also been observed by other authors [34, 35]. If there is a high number of old samples (generally > 60 years) in the control model, this can lead to a lower model slope, which would incorrectly assign negative EAA to young samples. This highlights the importance of having an age distribution in the control samples that matches that of the cases to be tested for differences in EAA.

Thus, we have shown that correcting for batch effects in the context of the epigenetic clock is important, especially when combining datasets from different sources for meta-analysis purposes. Batch effect correction is essential to remove technical variance that could affect the epigenetic age of the samples and confound biological interpretation.

### Sotos syndrome accelerates epigenetic ageing

Once we had corrected for potential batch effects in the data, we compared the epigenetic age acceleration (EAA) distributions between each of the developmental disorders studied and our control set. For a given sample, a positive EAA indicates that the epigenetic (biological) age of the sample is higher than the one expected for someone with that chronological age. In other words, it means that the epigenome of that person resembles the epigenome of an older individual. The opposite is true when a negative EAA is found (i.e. the epigenome looks younger than expected).

For the main screen, we selected those control samples with the same age range as the one present when aggregating all the cases (0 to 55 years), since this permits the development of a common control (background) model and to compare the statistical significance of the results across developmental disorders. Only those developmental disorders that satisfied our filtering criteria were considered for the screen (at least 5 samples available for the developmental disorder, with 2 of them presenting a chronological age ≥ 20 years, Fig. 1a, Table 1 and Additional file 2). Given that the blood composition changes with age (changes in the different cell types proportions, which can affect bulk DNA methylation measurements), we used models with and without cell composition correction (CCC), correcting for batch effects in both of them (see Methods). It is important to mention that EAA_with CCC_ is conceptually similar to the previously reported measure of ‘intrinsic EAA’ (IEAA) [18, 36].

The results from the screen are portrayed in Fig. 2a. Most syndromes do not show evidence of accelerated epigenetic ageing, but Sotos syndrome presents a clear positive EAA (median EAA_with CCC_ = + 7.64 years, median EAA_without CCC_ = + 7.16 years), with p-values considerably below the significance level of 0.01 after Bonferroni correction (p-value_corrected, with CCC_ = 3.40 · 10^−9^, p-value_corrected, without CCC_ = 2.61 · 10^−7^). Additionally, Rett syndrome (median EAA_with CCC_ = + 2.68 years, median EAA_without CCC_ = + 2.46 years, p-value_corrected, with CCC_ = 0.0069, p-value_corrected, without CCC_ = 0.0251) and Kabuki syndrome (median EAA_with CCC_ = − 1.78 years, median EAA_without_ CCC = − 2.25 years, p-value_corrected, with CCC_ = 0.0011, p-value_corrected, without CCC_ = 0.0035) reach significance, with a positive and negative EAA respectively. Finally, fragile × syndrome (FXS) shows a positive EAA trend (median EAA_with CCC_ = + 2.44 years, median EAA_without CCC_ = + 2.88 years) that does not reach significance in our screen (p-value_corrected, with CCC_ = 0.0680, p-valuecorrected, without CCC = 0.0693).

**Fig. 2.**
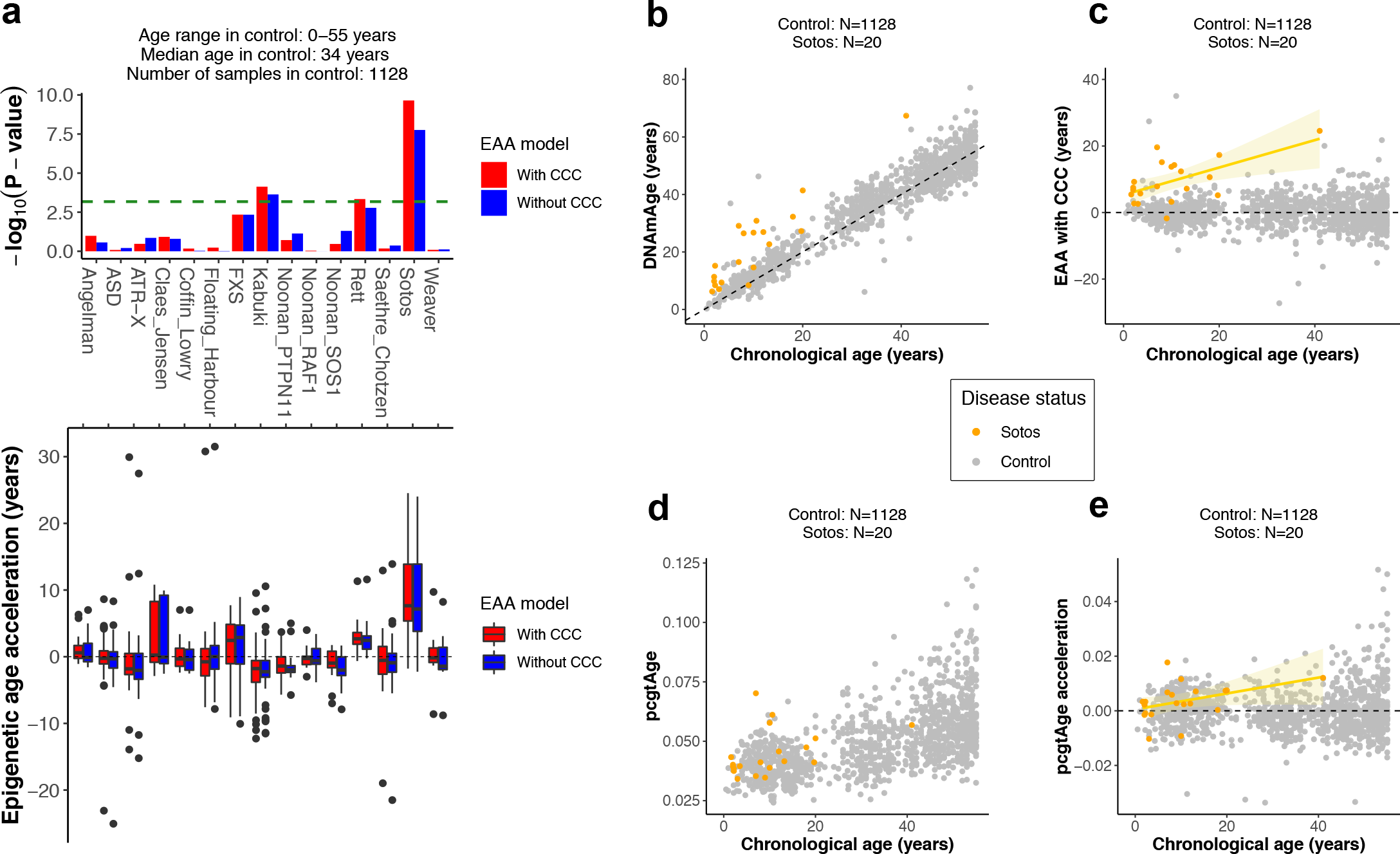
Sotos syndrome accelerates epigenetic ageing. **a.** Screening for epigenetic age acceleration (EAA) in developmental disorders. The upper panel shows the p-values derived from comparing the EAA distributions for the samples in a given developmental disorder and the control (two-sided Wilcoxon’s test). The dashed green line displays the significance level of α = 0.01 after Bonferroni correction. The bars above the green line reach statistical significance. The lower panel displays the actual EAA distributions, which allows assessing the direction of the EAA (positive or negative). In red: EAA model with cell composition correction (CCC). In blue: EAA model without CCC. ASD: autism spectrum disorder. ATR-X: alpha thalassemia/mental retardation X-linked syndrome. FXS: fragile × syndrome. **b.** Scatterplot showing the relation between epigenetic age (*DNAmAge*) according to Horvath’s model [8] and chronological age of the samples for Sotos (orange) and control (grey). Each sample is represented by one point. The black dashed line represents the diagonal to aid visualisation. **c.** Scatterplot showing the relation between the epigenetic age acceleration (EAA) and chronological age of the samples for Sotos (orange) and control (grey). Each sample is represented by one point. The yellow line represents the linear model *EAA ~ Age*, with the standard error shown in the light yellow shade. **d.** Scatterplot showing the relation between the score for the epigenetic mitotic clock (*pcgtAge*) [37] and chronological age of the samples for Sotos (orange) and control (grey). Each sample is represented by one point. A higher value of *pcgtAge* is associated with a higher number of cell divisions in the tissue. **e.** Scatterplot showing the relation between the epigenetic mitotic clock (*pcgtAge*) acceleration and chronological age of the samples for Sotos (orange) and control (grey). Each sample is represented by one point. The yellow line represents the linear model *pcgtAge*_*acceleration*_ ~ *Age*, with the standard error shown in the light yellow shade.

Next, we tested the effect of changing the median age used to build the healthy control model (i.e. the median age of the controls) on the screening results (Additional file 1: Figure S2A). Sotos syndrome is robust to these changes, whilst Rett, Kabuki and FXS are much more sensitive to the control model used. This again highlights the importance of choosing an appropriate age-matched control when testing for epigenetic age acceleration, given that Horvath’s epigenetic clock underestimates epigenetic age for advanced chronological ages [34, 35].

Moreover, all but one of the Sotos syndrome patients (19/20 = 95%) show a consistent deviation in EAA (with CCC) in the same direction (Fig. 2b, c), which is not the case for the rest of the disorders, with the exception of Rett syndrome (Additional file 1: Figure S2B). Even though the data suggest that there are already some methylomic changes at birth, the EAA seems to increase with age in the case of Sotos patients (Fig. 2c). This implies that at least some of the changes that normally affect the epigenome with age are happening at a faster rate in Sotos syndrome patients during their lifespan (as opposed to the idea that the Sotos epigenetic changes are only acquired during prenatal development and remain constant afterwards).

Finally, we investigated whether Sotos syndrome leads to a higher rate of (stem) cell division in blood when compared with our healthy population. We used a reported epigenetic mitotic clock (*pcgtAge*) that makes use of the fact that some CpGs in promoters that are bound by Polycomb group proteins become hypermethylated with age. This hypermethylation correlates with the number of cell divisions in the tissue and is also associated with an increase in cancer risk [37]. We found a trend suggesting that the epigenetic mitotic clock might be accelerated in Sotos patients (p-value = 0.0112, Fig. 2d, e), which could explain the higher cancer predisposition reported in these patients and might relate to their overgrowth [38].

Consequently, we report that people with Sotos syndrome present an accelerated epigenetic age, which makes their epigenome look, on average, more than 7 years older than expected. These changes seem to be the consequence of a higher ticking rate of the epigenetic clock (or at least part of its machinery), with epigenetic age acceleration increasing during lifespan: the youngest Sotos patient (1.6 years) has an EAA_with CCC_ = 5.43 years and the oldest (41 years) has an EAA_with CCC_ = 24.53 years. Additionally, Rett syndrome, Kabuki syndrome and fragile × syndrome could also have their epigenetic ages affected, but more evidence is required to be certain about this conclusion.

### Physiological ageing and Sotos syndrome share methylation changes and the genomic context in which they occur

Sotos syndrome is caused by loss-of-function heterozygous mutations in the *NSD1* gene, a histone H3K36 methyltransferase [39, 40]. These mutations lead to a specific DNA methylation signature in Sotos patients, potentially due to the crosstalk between the histone and DNA methylation machinery [40]. In order to gain a more detailed picture of the reported epigenetic age acceleration, we decided to compare the genome-wide (or at least array-wide) changes observed in the methylome during ageing with those observed in Sotos syndrome. For this purpose, we identified differentially methylated positions (DMPs) for both conditions (see Methods). Ageing DMPs (aDMPs), were composed almost equally of CpG sites that gain methylation with age (i.e. become hypermethylated, 51.69%) and CpG sites that lose methylation with age (i.e. become hypomethylated, 48.31%, barplot in Fig. 3a), a picture that resembles previous studies [41]. On the contrary, DMPs in Sotos were dominated by CpGs that decrease their methylation level in individuals with the syndrome (i.e. hypomethylated, 99.27%, barplot in Fig. 3a), consistent with previous reports [40].

**Fig. 3.**
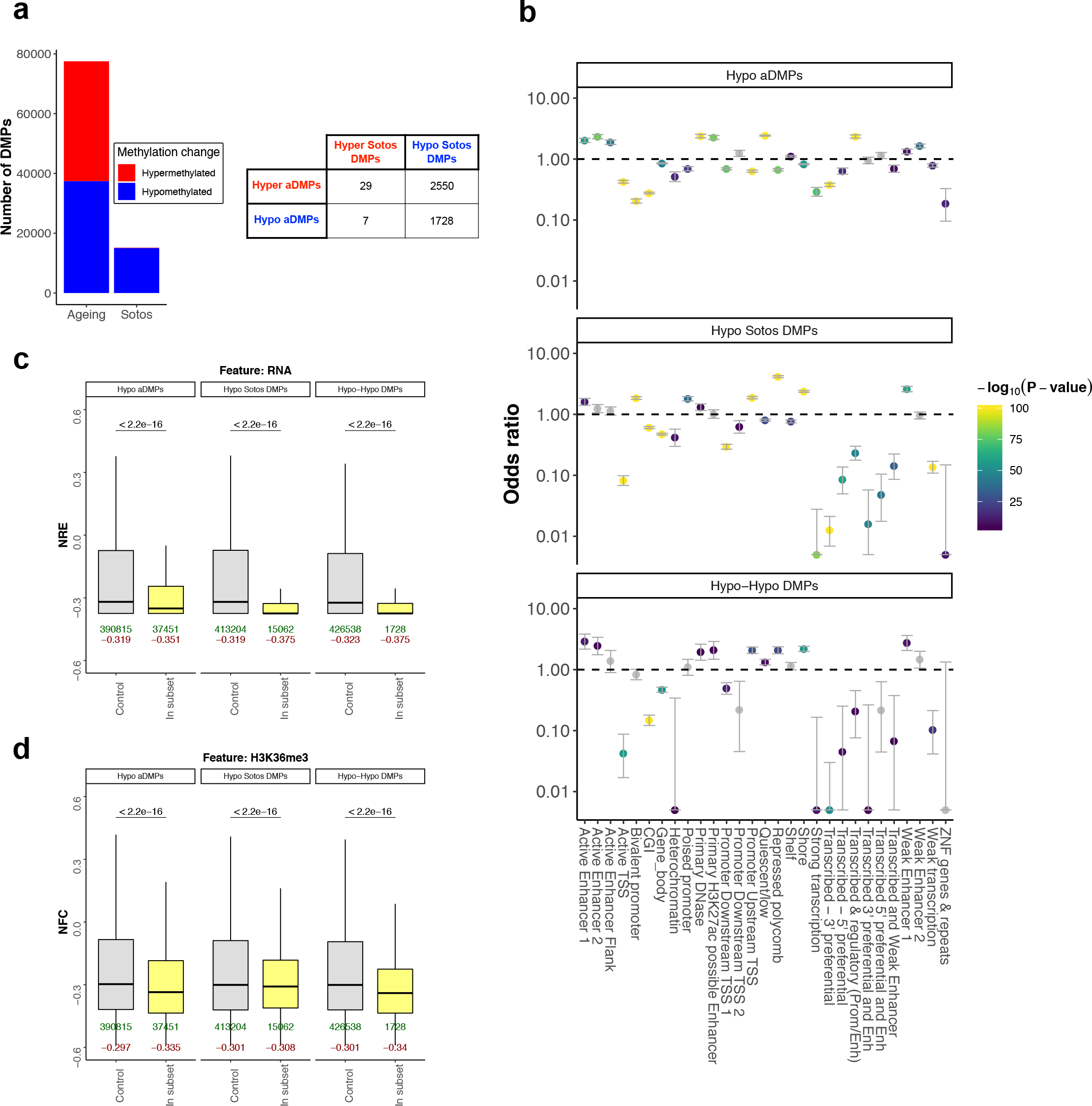
Comparison between the DNA methylation changes during physiological ageing and in Sotos. **a.** On the left: barplot showing the total number of differentially methylated positions (DMPs) found during physiological ageing and in Sotos syndrome. CpG sites that increase their methylation levels with age in our healthy population or those that are elevated in Sotos patients (when compared with a control) are displayed in red. Conversely, those CpG sites that decrease their methylation levels are displayed in blue. On the right: table that represents the intersection between the ageing (aDMPs) and the Sotos DMPs. The subset resulting from the intersection between the hypomethylated DMPs in ageing and Sotos is called the ‘Hypo-Hypo DMPs’ subset (N=1728). **b.** Enrichment for the categorical (epi)genomic features considered when comparing the different genome-wide subsets of differentially methylated positions (DMPs) in ageing and Sotos against a control (see Methods). The y-axis represents the odds ratio (OR), the error bars show the 95% confidence interval for the OR estimate and the colour of the points codes for −log_10_(p-value) obtained after testing for enrichment using Fisher’s exact test. An OR > 1 shows that the given feature is enriched in the subset of DMPs considered, whilst an OR < 1 shows that it is found less than expected. In grey: features that did not reach significance using a significance level of α = 0.01 after Bonferroni correction. **c.** Boxplots showing the distributions of the ‘normalised RNA expression’ (NRE) when comparing the different genome-wide subsets of differentially methylated positions (DMPs) in ageing and Sotos against a control (see Methods). NRE represents normalised mean transcript abundance in a window of ± 200 bp from the CpG site coordinate (DMP) being considered (see Methods). The p-values (two-sided Wilcoxon’s test, before multiple testing correction) are shown above the boxplots. The number of DMPs belonging to each subset (in green) and the median value of the feature score (in dark red) are shown below the boxplots. **d.** Same as c., but showing the ‘normalised fold change’ (NFC) for the H3K36me3 histone modification (representing normalised mean ChIP-seq fold change for H3K36me3 in a window of ± 200 bp from the DMP being considered, see Methods).

Then, we compared the intersections between the hypermethylated and hypomethylated DMPs in ageing and Sotos. Most of the DMPs were specific for ageing or Sotos (i.e. they did not overlap), but a subset of them were shared (table in Fig. 3a). Interestingly, there were 1728 DMPs that became hypomethylated both during ageing and in Sotos (‘Hypo-Hypo DMPs’). This subset of DMPs is of special interest because it could be used to understand in more depth some of the mechanisms that drive hypomethylation during physiological ageing. Thus, we tested whether the different subsets of DMPs are found in specific genomic contexts (Additional file 1: Figures S3A,B). DMPs that are hypomethylated during ageing and in Sotos were both enriched (odds ratio >1) in enhancer categories (such as ‘active enhancer 1’ or ‘weak enhancer 1’, see the chromatin state model used, from the K562 cell line, in Methods) and depleted (odds ratio <1) for active transcription categories (such as ‘active TSS’ or ‘strong transcription’), which was also observed in the ‘Hypo-Hypo DMPs’ subset (Fig. 3b). Interestingly, age-related hypomethylation in enhancers seems to be a characteristic of both humans [42, 43] and mice [24]. Furthermore, both *de novo* DNA methyltransferases (DNMT3A and DNMT3B) have been shown to bind in an H3K36me3-dependent manner to active enhancers [44], consistent with our results.

When looking at the levels of total RNA expression (depleted for rRNA) in blood, we confirmed a significant reduction in the RNA levels around these hypomethylated DMPs when compared with the controls sets (Fig. 3c, see Methods for more details on how the control sets were defined). Interestingly, hypomethylated DMPs in both ageing and Sotos were depleted from gene bodies (Fig. 3b) and were located in areas with lower levels of H3K36me3 when compared with the control sets (Fig. 3d, Additional file 1: Figure S3B). Moreover, hypomethylated aDMPs and hypomethylated Sotos DMPs where both generally enriched or depleted for the same histone marks in blood (Additional file 1: Figure S3B), which adds weight to the hypothesis that they share the same genomic context and could become hypomethylated through similar molecular mechanisms.

Intriguingly, we also identified a subset of DMPs (2550) that were hypermethylated during ageing and hypomethylated in Sotos (Fig. 3a). These ‘Hyper-Hypo DMPs’ seem to be enriched for categories such as ‘bivalent promoter’ and ‘repressed polycomb’ (Additional file 1: Figure S3A), which are normally associated with developmental genes [45, 46]. These categories are also a defining characteristic of the hypermethylated aDMPs, highlighting that even though the direction of the DNA methylation changes is different in some ageing and Sotos DMPs, the genomic context in which they happen is shared.

Finally, we looked at the DNA methylation patterns in the 353 epigenetic clock CpG sites for the Sotos samples. For each clock CpG site, we modelled the changes of DNA methylation during the lifespan in the healthy control individuals and then calculated the deviations from these patterns for the Sotos samples (Additional file 1: Figure S3C, see Methods). As expected, the landscape of clock CpG sites is dominated by hypomethylation in the Sotos samples, although only a small fraction of the clock CpG sites seems to be significantly affected (Additional file 1: Figure S3D). Overall, we confirmed the trends reported for the genome-wide analysis (Additional file 1: Figures S3E-G). However, given the much smaller number of CpG sites to consider in this analysis, very few comparisons reached significance.

We have demonstrated that the ageing process and Sotos syndrome share a subset of hypomethylated CpG sites that is characterised by an enrichment in enhancer features and a depletion of active transcription activity. This highlights the usefulness of developmental disorders as a model to study the mechanisms that may drive the changes in the methylome with age, since they permit stratification of the ageing DMPs into different functional categories that are associated with alterations in the function of specific genes and hence specific molecular components of the epigenetic ageing clock.

### Sotos syndrome is associated with a decrease of methylation Shannon entropy in the epigenetic clock CpG sites

Shannon entropy can be used in the context of DNA methylation analysis to estimate the information content stored in a given set of CpG sites. Shannon entropy is minimised when the methylation levels of all the CpG sites are either 0% or 100% and maximised when all of them are 50% (see Methods). Previous reports have shown that the Shannon entropy associated with the methylome increases with age, which implies that the epigenome loses information content [9, 12, 42]. We confirmed this genome-wide effect (i.e. considering all the CpG sites that passed our pre-processing pipeline) in our healthy samples, where we observed a positive Spearman correlation coefficient between chronological age and genome-wide Shannon entropy of 0.3984 (p-value = 3.21 · 10^−44^). This result was robust when removing outlier batches (Additional file 1: Figure S4C). Next, we tested whether Sotos patients present genome-wide Shannon entropy acceleration i.e. deviations from the expected genome-wide Shannon entropy for their age (see Methods). Despite detailed analysis, we did not find evidence that this was the case when looking genome-wide (p-value = 0.71, Fig. 4a, b, Additional file 1: Figure S4A).

**Fig. 4.**
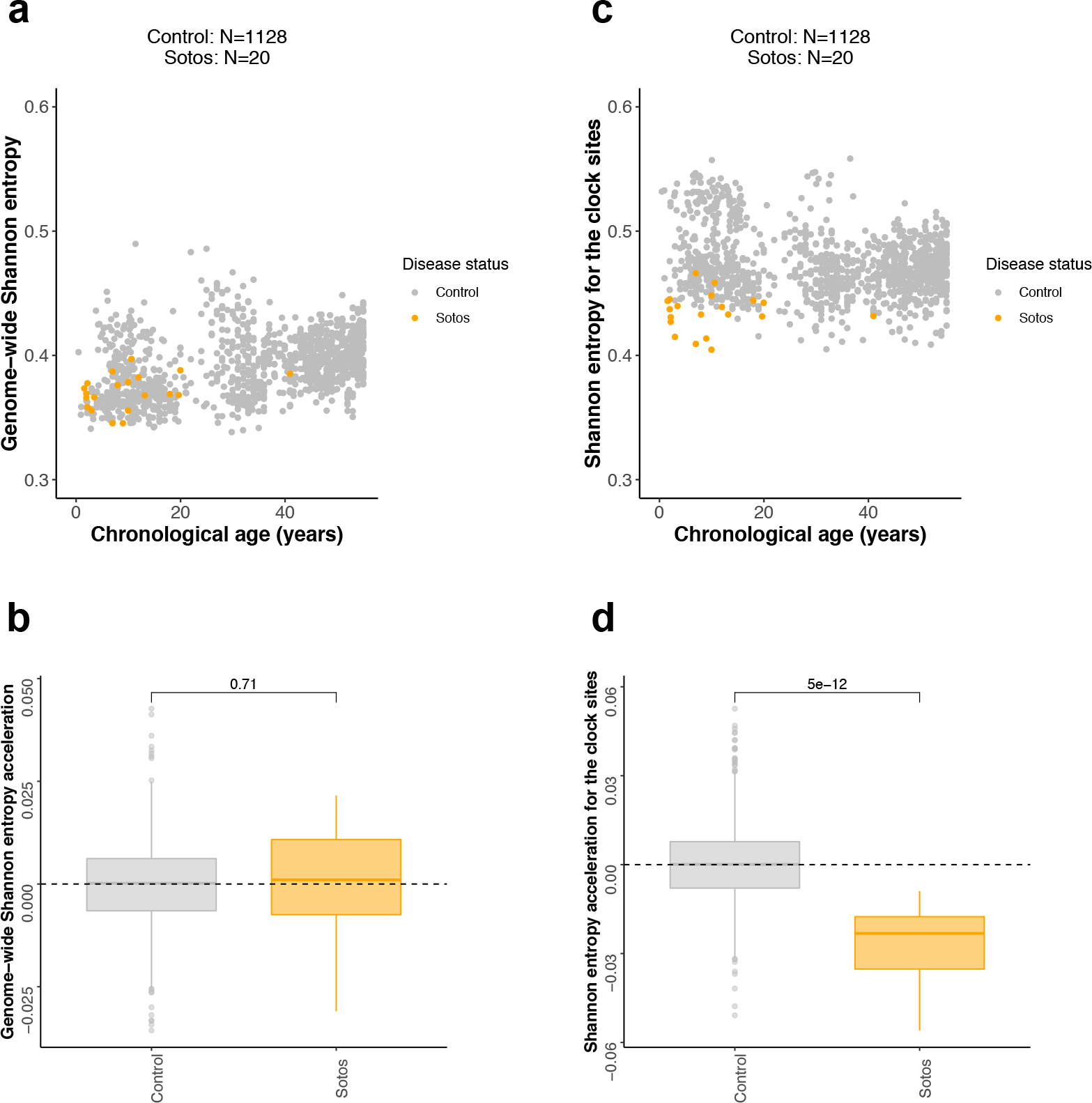
Analysis of methylation Shannon entropy during physiological ageing and in Sotos syndrome. **a.** Scatterplot showing the relation between genome-wide Shannon entropy (i.e. calculated using the methylation levels of all the CpG sites in the array) and chronological age of the samples for Sotos (orange) and healthy controls (grey). Each sample is represented by one point. **b.** Boxplots showing the distributions of genome-wide Shannon entropy acceleration (i.e. deviations from the expected genome-wide Shannon entropy for their age) for the control and Sotos samples. The p-value displayed on top of the boxplots was derived from a two-sided Wilcoxon’s test. **c.** Same as a., but using the Shannon entropy calculated only for the 353 CpG sites in the Horvath epigenetic clock. **d.** Same as b., but using the Shannon entropy calculated only for the 353 CpG sites in the Horvath epigenetic clock.

When we considered only the 353 clock CpG sites for the entropy calculations, the picture was different. Shannon entropy for the 353 clock sites slightly decreased with age in the controls when we included all the batches, showing the opposite direction when compared with the genome-wide entropy (Spearman correlation coefficient = −0.1223, p-value = 3.8166

10^−5^, Fig. 4c). However, when we removed the ‘Europe’ batch (which was an outlier even after pre-processing, Additional file 1: Figure S4D), this trend was reversed and we observed a weak increase of clock Shannon entropy with age (Spearman correlation coefficient = 0.1048, p-value = 8.6245 · 10^−5^). This shows that Shannon entropy calculations are very sensitive to batch effects, especially when considering a small number of CpG sites, and the results must be interpreted carefully.

Interestingly, the mean Shannon entropy across all the control samples was higher in the epigenetic clock sites (mean = 0.4726, Fig. 4c) with respect to the genome-wide entropy (mean = 0.3913, Fig. 4a). Sotos syndrome patients displayed a lower clock Shannon entropy when compared with the control (p-value = 5.0449 · 10^−12^, Fig. 4d, Additional file 1: Figure S4B), which is probably driven by the hypomethylation of the clock CpG sites. Furthermore, this highlights that the Horvath clock sites could have slightly different characteristics in terms of the methylation entropy associated with them when compared with the genome as a whole, something that to our knowledge has not been reported before.

## DISCUSSION

The epigenetic ageing clock has emerged as the most accurate biomarker of the ageing process and it seems to be a conserved property in mammalian genomes [5, 6]. However, we do not know yet whether the age-related DNA methylation changes measured are functional at all or whether they are related to some fundamental process of the biology of ageing. Developmental disorders in humans represent an interesting framework to look at the biological effects of mutations in genes that are fundamental for the integrity of the epigenetic landscape and other core processes, such as growth or neurodevelopment [29, 30]. Therefore, using a reverse genetics approach, we aimed to identify genes that disrupt aspects of the behaviour of the epigenetic ageing clock in humans.

Most of the studies have looked at the epigenetic ageing clock using Horvath’s model [8], which has a ready-to-use online calculator for epigenetic age [47]. This has clearly simplified the computational process and helped a lot of research groups to test the behaviour of the epigenetic clock in their system of interest. However, this has also led to the treatment of the epigenetic clock as a ‘black-box’, without critical assessment of the statistical methodology behind it. Therefore, we decided to benchmark the main steps involved when estimating epigenetic age acceleration (pre-processing of the raw data from methylation arrays and cell composition deconvolution algorithms), to quantify the effects of technical variation on the epigenetic clock predictions and to assess the impact of the control age distribution on the epigenetic age acceleration calculations. Previous attempts to account for technical variation have used the first 5 principal components (PCs) estimated directly from the DNA methylation data [23]. However, this approach potentially removes meaningful biological variation. For the first time, we have shown that it is possible to use the control probes from the 450K array to readily correct for batch effects in the context of the epigenetic clock, which reduces the error associated with the predictions and decreases the likelihood of reporting a false positive. Furthermore, we have confirmed the suspicion that Horvath’s model underestimates epigenetic age for older ages [34, 35] and assessed the impact of this bias in the screen for epigenetic age acceleration.

The results from our screen strongly suggest that Sotos syndrome accelerates epigenetic ageing. Sotos syndrome is caused by loss-of-function mutations in the *NSD1* gene [39, 40], which encodes a histone H3 lysine 36 (H3K36) methyltransferase. This leads to a phenotype which can include pre-natal and post-natal overgrowth, facial gestalt, advanced bone age, developmental delay, higher cancer predisposition and, in some cases, heart defects [38]. Remarkably, many of these characteristics could be interpreted as ageing-like, identifying Sotos syndrome as a potential human model of accelerated physiological ageing.

NSD1 catalyses the addition of either monomethyl (H3K36me) or dimethyl groups (H3K36me2) and indirectly regulates the levels of trimethylation (H3K36me3) by altering the availability of the monomethyl and dimethyl substrates for the trimethylation enzymes (SETD2 in humans, whose mutations cause a ‘Sotos-like’ overgrowth syndrome) [48, 49]. H3K36 methylation has a complex role in the regulation of transcription [48] and has been shown to regulate nutrient stress response in yeast [50]. Moreover, experiments in model organisms (yeast and worm) have demonstrated that mutations in H3K36 methyltranferases decrease lifespan and, remarkably, mutations in H3K36 demethylases increase it [51–53].

In humans, DNA methylation patterns are established and maintained by three conserved enzymes: the maintenance DNA methyltransferase DNMT1 and the *de novo* DNA methyltransferases DNMT3A and DNMT3B [54]. Both DNMT3A and DNMT3B contain PWWP domains that can read the H3K36me3 histone mark [55, 56]. Therefore, the H3K36 methylation landscape can influence DNA methylation levels in specific genomic regions through the recruitment of the *de novo* DNA methyltransferases. Mutations in the PWWP domain of DNMT3A impair its binding to H3K36me2 and H3K36me3 and cause an undergrowth disorder in humans (microcephalic dwarfism) [57]. This redirects DNMT3A, which is normally targeted to H3K36me2 and H3K36me3 throughout the genome, to DNA methylation valleys (DMVs, aka DNA methylation canyons), which become hypermethylated [57]; a phenomenon that also seems to happen during physiological ageing in humans [42, 58, 59] and mice [24]. DMVs are hypomethylated domains conserved across cell types and species, often associated with Polycomb-regulated developmental genes and marked by bivalent chromatin (with H3K27me3 and H3K4me3) [60–63]. Therefore, we suggest a model (Fig. 5) where the reduction in the levels of H3K36me2 and/or H3K36me3, caused by a proposed decrease in H3K36 methylation maintenance during ageing or NSD1 function in Sotos syndrome, could lead to hypomethylation in many genomic regions (because DNMT3A is recruited less efficiently) and hypermethylation in DMVs (because of the higher availability of DNMT3A). Indeed, we observe enrichment for categories such as ‘bivalent promoter’ or ‘repressed polycomb’ in the hypermethylated DMPs in Sotos and ageing (Additional file 1: Figure S3A), which is also supported by higher levels of Polycomb Repressing Complex 2 (PRC2, represented by EZH2) and H3K27me3, the mark deposited by PRC2 (Additional file 1: Figure S3B). This is also consistent with the results obtained for the epigenetic mitotic clock [37], where we observe a trend towards increased hypermethylation of Polycomb-bound regions in Sotos patients.

**Fig. 5.**
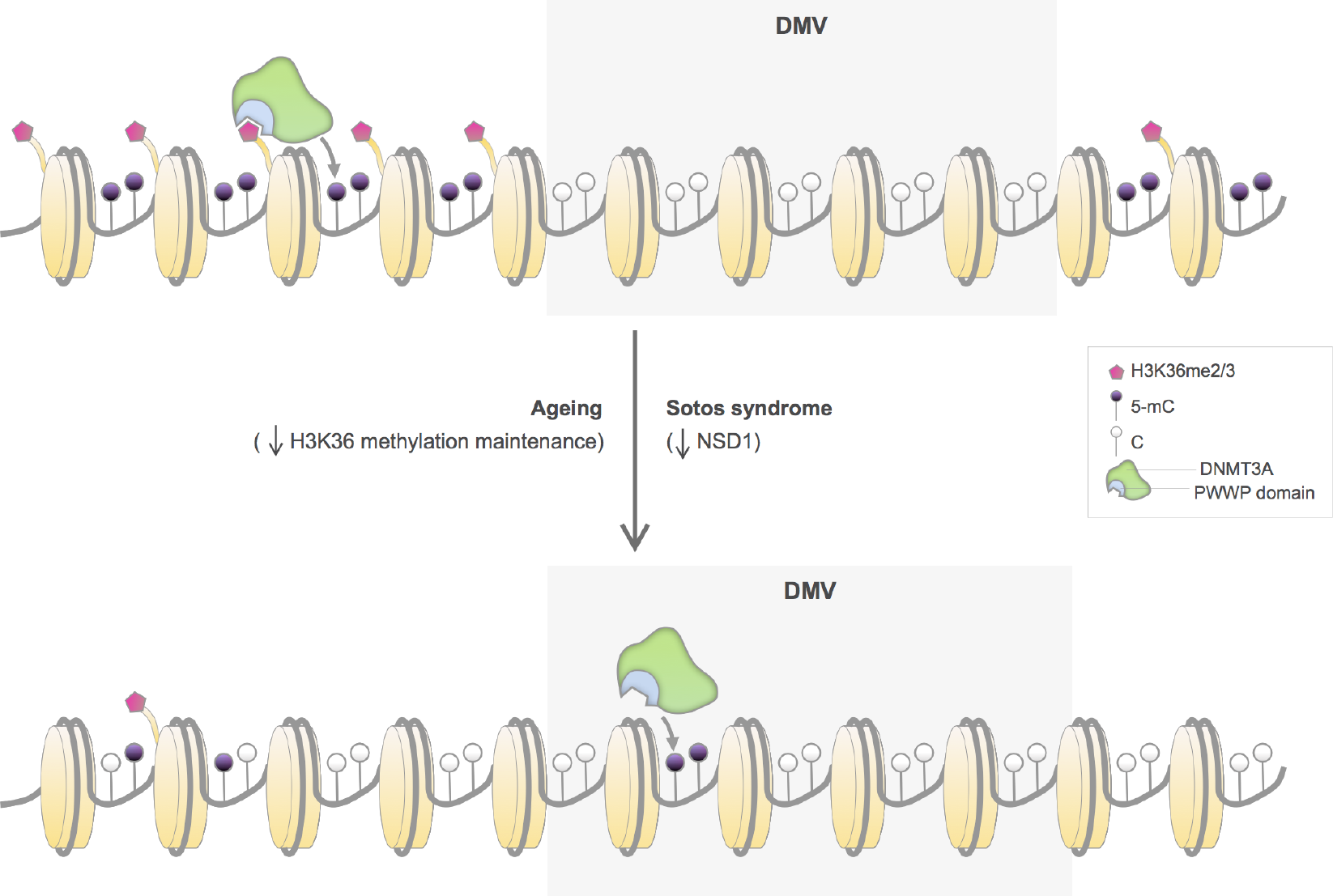
Proposed model that highlights the role of H3K36 methylation maintenance on epigenetic ageing. The H3K36me2/3 mark allows recruiting *de novo* DNA methyltransferases DNMT3A (in green) and DNMT3B (not shown) through their PWWP domain (in blue) to different genomic regions (such as gene bodies or pericentric heterochromatin) [56, 94, 95], which leads to the methylation of the cytosines in the DNA of these regions (5-mC, black lollipops). On the contrary, DNA methylation valleys (DMVs) are conserved genomic regions that are normally found hypomethylated and associated with Polycomb-regulated developmental genes [60–63]. During ageing, the H3K36 methylation machinery could become less efficient at maintaining the H3K36me2/3 landscape. This would lead to a relocation of *de novo* DNA methyltransferases from their original genomic reservoirs (which would become hypomethylated) to other non-specific regions such as DMVs (which would become hypermethylated and potentially lose their normal boundaries), with functional consequences for the tissues. This is also partially observed in patients with Sotos syndrome, where mutations in NSD1 potentially affect H3K36me2/3 patterns and accelerate the epigenetic ageing clock as measured with the Horvath model [8]. Given that DNMT3B is enriched in the gene bodies of highly transcribed genes [56] and that we found these regions depleted in our differential methylation analysis, we hypothesise that the hypermethylation of DMVs could be mainly driven by DNMT3A instead. However, it is important to mention that our analysis does not discard a role of DNMT3B during epigenetic ageing.

A recent preprint has shown that loss-of-function mutations in DNMT3A, which cause Tatton-Brown-Rahman overgrowth syndrome, also lead to a higher ticking rate of the epigenetic ageing clock [64]. They also report positive epigenetic age acceleration in Sotos syndrome and negative acceleration in Kabuki syndrome, consistent with our results. Furthermore, they observe a DNA methylation signature in the DNMT3A mutants characterised by widespread hypomethylation, with a modest enrichment of DMPs in regions upstream of the transcription start site, shores and enhancers [64], which we also detect in our ‘Hypo-Hypo DMPs’ (those that become hypomethylated both during physiological ageing and in Sotos). Therefore, the hypomethylation observed in our ‘Hypo-Hypo DMPs’ is consistent with a reduced methylation activity of DNMT3A, which in our system could be a consequence of the decreased recruitment of DNMT3A to genomic regions that have lost H3K36 methylation (Fig. 5).

Interestingly, H3K36me3 is required for the selective binding of the *de novo* DNA methyltransferase DNMT3B to the bodies of highly transcribed genes [56]. Furthermore, DNMT3B loss reduces gene-body methylation, which leads to intragenic spurious transcription (aka cryptic transcription) [65]. An increase in this so-called cryptic transcription seems to be a conserved feature of the ageing process [52]. Therefore, the changes observed in the ‘Hypo-Hypo DMPs’ could theoretically be a consequence of the loss of H3K36me3 and the concomitant inability of DNMT3B to be recruited to gene bodies. However, the ‘Hypo-Hypo DMPs’ were depleted for H3K36me3, active transcription and gene bodies when compared with the rest of the probes in the array (Fig. 3b-d), prompting us to suggest that the DNA methylation changes observed are likely mediated by DNMT3A instead (Fig. 5). Nevertheless, it is worth mentioning that the different biological replicates for the blood H3K36me3 ChIP-seq datasets were quite heterogeneous and that the absolute difference in the case of the hypomethylated Sotos DMPs, although significant due to the big sample sizes, is quite small. Thus, we cannot exclude the existence of this mechanism during human ageing and an exhaustive study on the prevalence of cryptic transcription in humans and its relation to the ageing methylome should be carried out.

Because of the way that the Horvath epigenetic clock was trained [8], it is likely that its constituent 353 CpG sites are a low-dimensional representation of the different genome-wide processes that are eroding the epigenome with age. Our analysis has shown that these 353 CpG sites are characterised by a higher Shannon entropy when compared with the rest of the genome, which is dramatically decreased in the case of Sotos patients. This could be related to the fact that the clock CpGs are enriched in regions of bivalent chromatin (marked by H3K27me3 and H3K4me3), conferring a more dynamic or plastic regulatory state with levels of DNA methylation deviated from the collapsed states of 0 or 1. Interestingly, EZH2 (part of polycomb repressing complex 2, responsible for H3K27 methylation) is an interacting partner of DNMT3A and NSD1, with mutations in NSD1 affecting the genome-wide levels of H3K27me3 [66]. Furthermore, Kabuki syndrome was weakly identified in our screen as having an epigenome younger than expected, which could be related to the fact that they show post-natal dwarfism [67, 68]. Kabuki syndrome is caused by loss-of-function mutations in KMT2D [67, 68], a major mammalian H3K4 mono-methyltransferase [69]. Additionally, H3K27me3 and H3K4me3 levels can affect lifespan in model organisms [3]. It will be interesting to test whether bivalent chromatin is a general feature of multi-tissue epigenetic ageing clocks.

Thus, DNMT3A, NSD1 and the machinery in control of bivalent chromatin (such as EZH2 and KMT2D) contribute to an emerging picture on how the mammalian epigenome is regulated during ageing, which could open new avenues for anti-ageing drug development. Mutations in these proteins lead to different developmental disorders with impaired growth defects [29], with DNMT3A, NSD1 and potentially KMT2D also affecting epigenetic ageing. Interestingly, EZH2 mutations (which cause Weaver syndrome, Table 1) do not seem to affect the epigenetic clock in our screen. However, this syndrome has the smallest number of samples (7) and this could limit the power to detect any changes.

Our screen has also revealed that Rett syndrome and fragile × syndrome (FXS) could potentially have an accelerated epigenetic age. It is worth noting that FXS is caused by an expansion of the CGG trinucleotide repeat located in the 5’ UTR of the *FMR1* gene [70]. Interestingly, Huntington’s disease, caused by a trinucleotide repeat expansion of CAG, has also been shown to accelerate epigenetic ageing of human brain [23], pointing towards trinucleotide repeat instability as an interesting molecular mechanism to look at from an ageing perspective. It is important to notice that the conclusions for Rett syndrome, FXS and Kabuki syndrome were very dependent on the age range used in the healthy control (Additional file 1: Figure S2A) and these results must therefore be treated with caution.

Our study has several limitations that we tried to address in the best possible way. First of all, given that DNA methylation data for patients with developmental disorders is relatively rare, some of the sample sizes were quite small. It is thus possible that some of the other developmental disorders assessed are epigenetically accelerated but we lack the power to detect this. Furthermore, people with the disorders tend to get sampled when they are young i.e. before reproductive age. Horvath’s clock adjusts for the different rates of change in the DNA methylation levels of the clock CpGs before and after reproductive age (20 years in humans) [8], but this could still have an effect on the predictions, especially if the control is not properly age-matched. Our solution was to discard those developmental disorders with less than 5 samples and we required them to have at least 2 samples with an age ≥ 20 years, which reduced the list of final disorders included to the ones listed in Table 1.

Future studies should increase the sample size and follow the patients during their entire lifespan in order to confirm our findings. Furthermore, it would be interesting to identify mutations that affect, besides the mean, the variance of epigenetic age acceleration, since changes in methylation variability at single CpG sites with age have been associated with fundamental ageing mechanisms [42]. Finally, testing the influence of H3K36 methylation on the epigenetic clock and lifespan in mice will provide deeper mechanistic insights.

## CONCLUSIONS

The epigenetic ageing clock has created a new methodological paradigm to study the ageing process in humans. However, the molecular mechanisms that control its ticking rate are still mysterious. In this study, by looking at patients with developmental disorders, we have demonstrated that Sotos syndrome accelerates epigenetic ageing and uncovered a potential role of the H3K36 methylation machinery as a key component of the *epigenetic maintenance system* in humans. We hope that this research will shed some light on the different processes that erode the human epigenetic landscape during ageing and provide a new hypothesis about the mechanisms behind the epigenetic ageing clock.

## METHODS

### Sample collection and annotation

We collected DNA methylation data generated with the Illumina Infinium HumanMethylation450 BeadChip (450K array) from human blood. In the case of the developmental disorder samples, we combined public data with data generated in-house for other clinical studies (Table 1, Additional file 2) [30]. We took all the data for developmental disorders that we could find in order to perform unbiased screening. The healthy samples used to build the control were mainly obtained from public sources (Additional file 3). Basic metadata (including the chronological age) was also stored. All the mutations in the developmental disorder samples were manually curated using Variant Effect Predictor [71] in the GRCh37 (hg19) human genome assembly. Those samples with a variant of unknown significance that had the characteristic DNA methylation signature of the disease were also included (they are labelled as ‘YES_predicted’ in Additional file 2). In the case of fragile × syndrome (FXS), only male samples with full mutation (>200 repeats) [70] were included in the final screen. As a consequence, only samples with a clear molecular and clinical diagnosis were kept for the final screen.

### Pre-processing, QC and filtering the data for the epigenetic clock calculations

Raw DNA methylation array data (IDAT files) were processed using the *minfi* R package [72]. Raw data were background-corrected using *noob* [73] before calculating the beta-values. In the case of the beta-values which are input to Horvath’s model, we observed that background correction did not have a major impact in the final predictions of epigenetic age acceleration in the control as long as we corrected for batch effects (Fig. 1c, Additional file 1: Figure S5A). We decided to keep the *noob* background correction step for consistency with the rest of the pipelines. Epigenetic age (*DNAmAge*) was calculated using the code from Horvath, which includes an internal normalisation step against a blood gold-standard [8]. The scripts are available in our GitHub repository [74] for the use of the community.

Quality control (QC) was performed in all samples. Following guidelines from the *minfi* package [72], only those samples that satisfied the following criteria were kept for the analysis: the sex predicted from the DNA methylation data was the same as the reported sex in the metadata, they passed BMIQ normalisation and 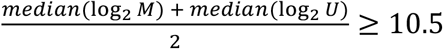, where *M* is the methylated intensity and *U* the unmethylated intensity for the array probes.

### Correcting for batch effects

In order to correct for batch effects that could confound the conclusions from our analysis, we decided to make use of the control probes available in the 450K array. These probes capture only technical variance in negative controls and different steps of the array protocol, such as bisulfite conversion, staining or hybridisation [32, 75]. We performed PCA (with centering but not scaling using the *prcomp* function in R) on the raw intensities of the control probes (847 probes · 2 channels = 1694 intensity values) for all our controls (N=2218) and cases (N=666) that passed QC (Fig. 1a). Including the technical PCs as covariates in the models to calculate epigenetic age acceleration (EAA) improved the error from the predictions in the controls (Fig. 1c, Additional file 1: Figure S5A). The optimal number of PCs was found by making use of the *findElbow* function from [76].

### Correcting for cell composition

The proportions of different blood cell types change with age and this can affect the methylation profiles of the samples. Therefore, when calculating epigenetic age acceleration, it is important to compare models with and without cell type proportions included as covariates [36]. Cell type proportions can be estimated from DNA methylation data using different deconvolution algorithms [77]. In the context of the epigenetic clock, most of the studies have used the Houseman method [78]. We have benchmarked different reference-based deconvolution strategies (combining different pre-processing steps, references and deconvolution algorithms) against a gold-standard dataset (GSE77797) [79]. Our results suggest that using the IDOL strategy [79] to build the blood reference (from the Reinius et al. dataset, GSE35069) [80], together with the Houseman algorithm [78] and some pre-processing steps (*noob* background correction, probe filtering, BMIQ normalisation), leads to the best cell type proportions estimates i.e. those that minimise the deviations between our estimates and the real cell type composition of the samples in the gold-standard dataset (Additional file 1: Figure S5B, Additional file 4). We used the *epidish* function from the *EpiDISH* R package [81] for these purposes.

### Calculating the epigenetic age acceleration (EAA) and performing the main screen

Only those developmental disorders for which we had at least 5 samples, with 2 of them with an age ≥ 20 years, were included in the main screen (N=367). Healthy samples that matched the age range of those disorders (0-55 years, N=1128) were used to train the following linear models (the *control models*):

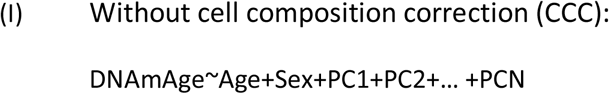

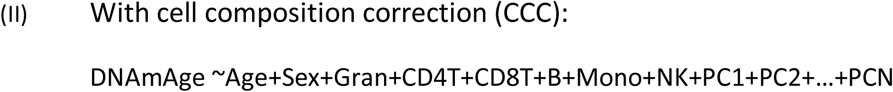

where *DNAmAge* is the epigenetic age calculated using Horvath’s model [8], *Age* is the chronological age, *PCN* is the *N*th technical PC obtained from the control probes (N=17 was the optimal, Fig. 1c) and *Gran*, *CD4T*, *CD8T*, *B*, *Mono* and *NK* are the different proportions of blood cell types as estimated with our deconvolution strategy. The linear models were fitted in R with the *lm* function, which uses least-squares.

The residuals from a control model represent the epigenetic age acceleration (EAA) for the different healthy samples, which should be centred around zero after batch effect correction (Additional file 1: Figure S1E, Fig. 1d). Then, the median absolute error (*MAE*) can be calculated as (Fig. 1c, Additional file 1: Figure S5A):

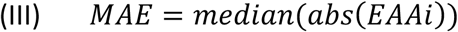

where *EAA*_*i*_ is the epigenetic age acceleration for a healthy sample from the control.

Once the control models are established, we can calculate the EAA for the different samples with a developmental disorder (cases) by taking the difference between the epigenetic age (*DNAmAge*) for the case sample and the predicted value from the corresponding control model (with or without cell composition correction). Finally, the distributions of the EAA for the different developmental disorders were compared against the EAA distribution for the healthy controls using a two-sided Wilcoxon’s test. P-values were adjusted for multiple testing using Bonferroni correction and a significance level of α = 0.01 was applied.

### Calculating *pcgtAge* and Shannon entropy

Raw DNA methylation data (IDAT files) was background-corrected using *noob* [73]. Next, we filtered out probes associated with SNPs, cross-reactive probes [82] and probes from the sex chromosomes, before performing BMIQ intra-array normalisation to correct for the bias in probe design [83]. Then, we calculated *pcgtAge* as the average of the beta-values for the probes that constitute the epigenetic mitotic clock [37]. It is worth noting that only 378 out of the 385 probes were left after our filtering criteria.

Shannon entropy was calculated as previously described [9]:

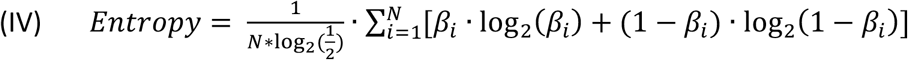

where β_i_ represents the methylation beta-value for the *i*th probe (CpG site) in the array, *N* = 428266 for the genome-wide entropy and *N* = 353 for Horvath clock sites entropy.

In order to calculate the *pcgtAge* and Shannon entropy acceleration, we followed a similar strategy to the one reported for EAA with CCC, fitting the following linear models:

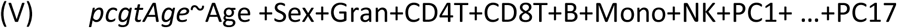

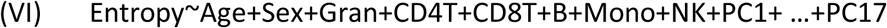

It is worth mentioning that we observed a remarkable effect of the batch on the Shannon entropy calculations, which generated high entropy variability for a given age (Additional file 1: Figure S4C,D). Thus, accounting for technical variation becomes crucial when assessing this type of data, even after background correction, probe filtering and BMIQ normalisation.

### Identifying differentially methylated positions (DMPs)

DMPs were identified using a modified version of the *dmpFinder* function in the *minfi* R package [72], where we accounted for other covariates. The ageing DMPs (aDMPs) were calculated using the control samples that were included in the screen (age range 0-55 years, N=1128) and the following linear model (p-values and regression coefficients were extracted for the *Age* covariate):

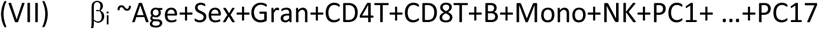

where β_i_ represents the methylation beta-value for the *i*th probe (CpG site) in the array.

The Sotos DMPs were calculated by comparing the Sotos samples (N=20) against the control samples (N=51) from the same dataset (GSE74432) [40] using the following linear model (p-values and regression coefficients were extracted for the *Disease_status* covariate):

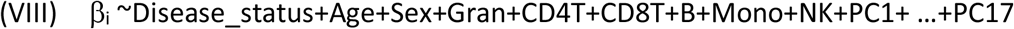

We selected as our final DMPs those CpG probes that survived our analysis after Bonferroni multiple testing correction with a significance level of α = 0.01.

### (Epi)genomic annotation of the CpG sites

Different (epi)genomic features were extracted for the CpG sites of interest. All the data were mapped to the *hg19* assembly of the human genome.

The continuous features were calculated by extracting the mean value in a window of ± 200 bp from the CpG site coordinate using the *pyBigWig* package [84]. We chose this window value based on the methylation correlation observed between neighbouring CpG sites in previous studies [85]. The continuous features included (Additional file 5):

- ChIP-seq data from ENCODE (histone modifications from peripheral blood mononuclear cells or PBMC; EZH2, as a marker of Polycomb Repressing Complex 2 binding, from B cells; RNF2, as a marker of Polycomb Repressing Complex 1 binding, from the K562 cell line). We obtained Z-scores (using the *scale* function in R) for the values of ‘fold change over control’ as calculated in ENCODE [86]. When needed, biological replicates of the same feature were aggregated by taking the mean of the Z-scores in order to obtain the ‘normalised fold change’ (NFC).
- ChIP-seq data for LaminB1 (GSM1289416, quantified as ‘normalised read counts’ or NRC) and Repli-seq data for replication timing (GSM923447, quantified as ‘wavelet-transformed signals’ or WTS). We used the same data from the IMR90 cell line as in [87].
- Total RNA-seq data (rRNA depleted, from PBMC) from ENCODE. We calculated Z-scores after aggregating the ‘signal of unique reads’ (*sur*) for both strands (+ and −) in the following manner:

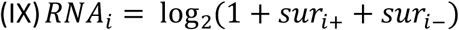

where *RNA*_*i*_ represents the RNA signal (that then needs to be scaled to obtain the ‘normalised RNA expression’ or NRE) for the *i*th CpG site.

The categorical features were obtained by looking at the overlap (using the *pybedtools* package) [88] of the CpG sites with the following:

- Gene bodies, from protein-coding genes as defined in the basic gene annotation of GENCODE release 29 [89].
- CpG islands (CGIs) were obtained from the UCSC Genome Browser [90]. Shores were defined as regions 0 to 2 kb away from CGIs in both directions and shelves as regions 2 to 4 kb away from CGIs in both directions as previously described [85, 91].
- Chromatin states were obtained from the K562 cell line in the Roadmap Epigenomics Project (based on imputed data, 25 states, 12 marks) [92]. A visualisation for the association between chromatin marks and chromatin states can be found in [93]. When needed for visualisation purposes, the 25 states were manually collapsed to a lower number of them.

We compared the different genomic features for each one of our subsets of CpG sites (hypomethylated aDMPs, hypomethylated Sotos DMPs, …) against a control set. This control set was composed of all the probes from the background set from which we removed the subset that we were testing. In the case of the comparisons against the 353 Horvath clock CpG sites, a background set of the 21368 (21K) CpG probes used to train the original Horvath model [8] was used. In the case of the genome-wide comparisons for ageing and Sotos syndrome, a background set containing all 428266 probes that passed our pre-processing pipeline (450K) was used.

The distributions of the scores from the continuous features were compared using a two-sided Wilcoxon’s test. In the case of the categorical features, we tested for enrichment using Fisher’s exact test.

### Differences in the clock CpGs beta-values for Sotos syndrome

To compare the beta-values of the Horvath clock CpG sites between our healthy samples and Sotos samples we fitted the following linear models in the healthy samples (*control CpG models*, Additional file 1: Figure S3C):

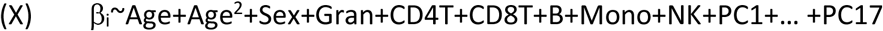

where β_i_ represents the methylation beta-values for the *i*th probe (CpG site) in the 353 CpG clock sites. The *Age*^*2*^ term allows accounting for non-linear relationships between chronological age and the beta-values.

Finally, we calculated the difference between the beta-values in Sotos samples and the predictions from the *control CpG models* and displayed these differences in an annotated heatmap (Additional file 1: Figure S3D).

### Code availability

All the code used to perform the analyses here presented can be found in our GitHub repository [74].

## LIST OF ABBREVIATIONS

aDMPs: differentially methylated positions during ageing
ASD: autism spectrum disorder
ATR-X: alpha thalassemia/mental retardation X-linked syndrome
CCC: cell composition correction
DMPs: differentially methylated positions
EAA: epigenetic age acceleration
FXS: fragile × syndrome
IEAA: intrinsic epigenetic age acceleration
iPSCs: induced pluripotent stem cells
MAE: median absolute error
PBMC: peripheral blood mononuclear cells
PCA: principal component analysis
PCs: principal components rRNA: ribosomal RNA
UTR: untranslated region
450K array: Illumina Infinium HumanMethylation450 BeadChip

## DECLARATIONS

### Ethics approval and consent to participate

The study protocol has been approved by the Western University Research Ethics Board (REB ID 106302), McMaster University and the Hamilton Integrated Research Ethics Boards (REB ID 13-653-T). All of the participants provided informed consent prior to sample collection. All of the samples and records were de-identified before any experimental or analytical procedures. The research was conducted in accordance with all relevant ethical regulations.

### Consent for publication

Not applicable.

### Availability of data and materials

Part of the DNA methylation data and metadata was obtained from the GEO public repository and are available under the following accession numbers: GSE104812, GSE111629, GSE116300, GSE35069 (to build the reference for cell composition estimation), GSE40279, GSE41273, GSE42861, GSE51032, GSE55491, GSE59065, GSE61496, GSE74432, GSE77797 (gold-standard for cell composition estimation), GSE81961 and GSE97362. The rest of the DNA methylation data and metadata (Europe, Feb_2016, Jun_2015, Mar_2014, May_2015, May_2016, Nov_2015, Oct_2014) are not publicly available at the time of the study as part of the conditions of the research ethical approval of the study, but may be available from the authors on reasonable request. All the code used to perform the analyses here presented can be found in the following GitHub repository [74].

## Competing interests

DEMH and TMS are founders and shareholders of Chronomics Limited, a UK-based company that provides epigenetic testing. WR is a consultant and shareholder of Cambridge Epigenetix.

## Funding

EMBL predoctoral fellowship (to DEMH); fellowship from the EMBL Interdisciplinary Postdoc (EI3POD) program under Marie Skłodowska-Curie Actions COFUND (grant number 664726; to MJB). WR acknowledges funding from BBSRC. JMT acknowledges funding from EMBL.

## Authors’ contributions

DEMH, TMS, WR and JMT designed the study. DEMH, EAE and MJB conducted data analysis. EAE and BS generated part of the DNA methylation dataset. MJB and OS provided crucial statistical input. DEMH, WR and JMT interpreted the data and wrote the manuscript. All authors read and approved the final manuscript.

## Acknowledgements

We thank all the members of the Reik and Thornton laboratories for helpful discussions. Specifically, we would like to acknowledge Melike Dönertas, Dr. Gos Micklem and Dr. Judith Zaugg for their advice during the last years. Furthermore, we thank all the authors and groups that answered our emails, allowed us to use the raw DNA methylation data that they generated and provided missing metadata; including Dr. Sanaa Choufani, Professor Rosanna Weksberg, Dr. Kimberly Aldinger, Dr. Sebastian Morán, Dr. Manuel Castro, Dr. Juan Sandoval, Dr. Pankaj Chopra, Dr. Akdes Serin-Harmanci, Dr. Weng Khong Lim, Dr. Reiner Schulz, Tina Wang, Dr. Barry Demchack and Professor Trey Ideker. Finally, we would like to thank Parvathi “Ale” Subbiah for designing the figure displaying our current working model.

