## Supplementary material for "Screening for genes that accelerate the epigenetic ageing clock in humans reveals a role for the H3K36 methyltransferase NSD1": Additional_file_1.docx

ADDITIONAL FILE 1: SUPPLEMENTARY FIGURES

**Figure S1**

**c**

**f**

**e**

**b**

**a**

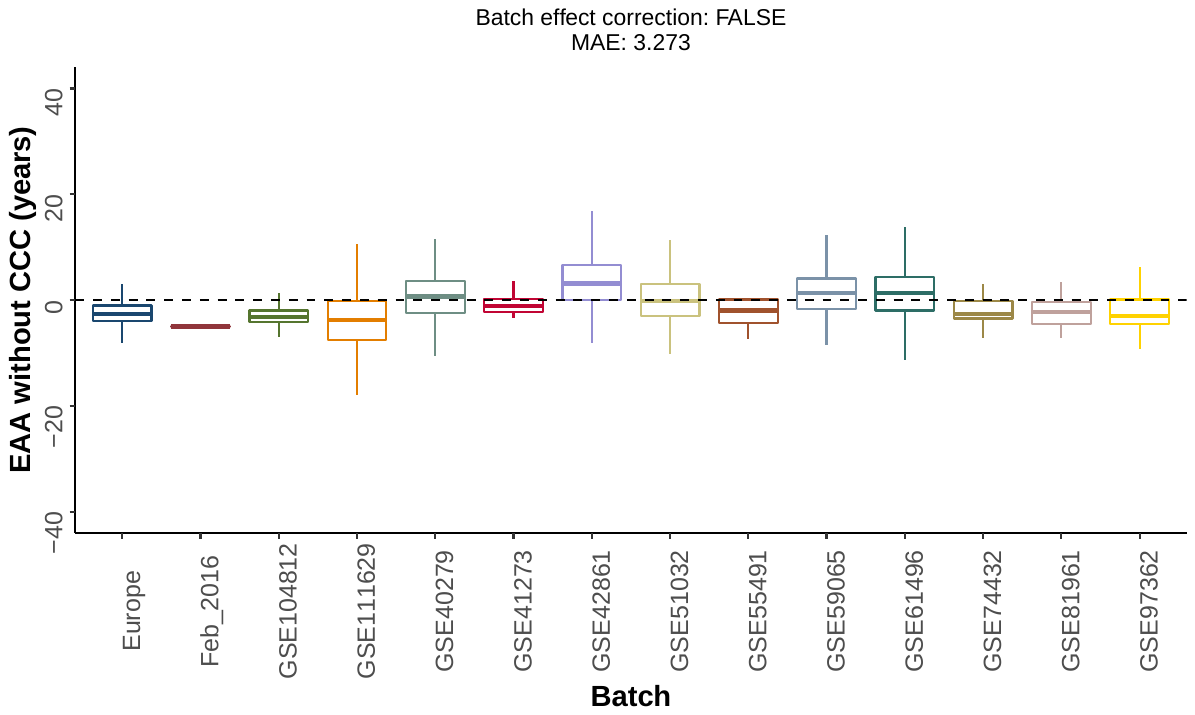

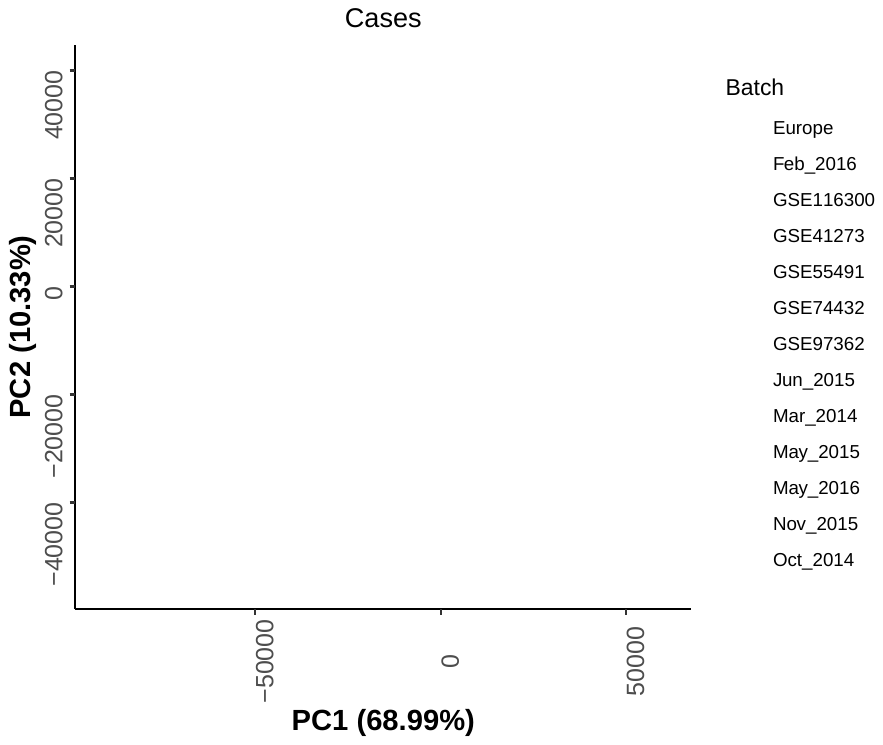

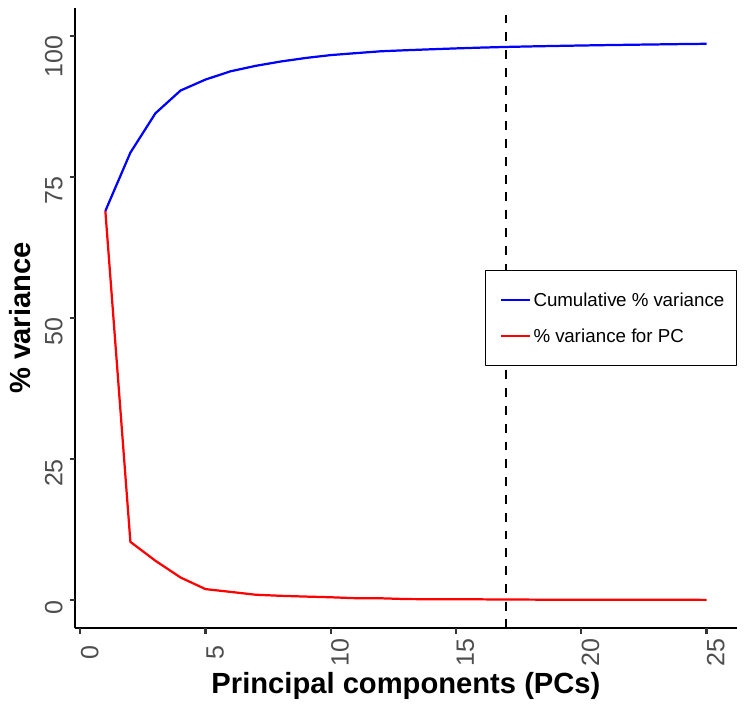

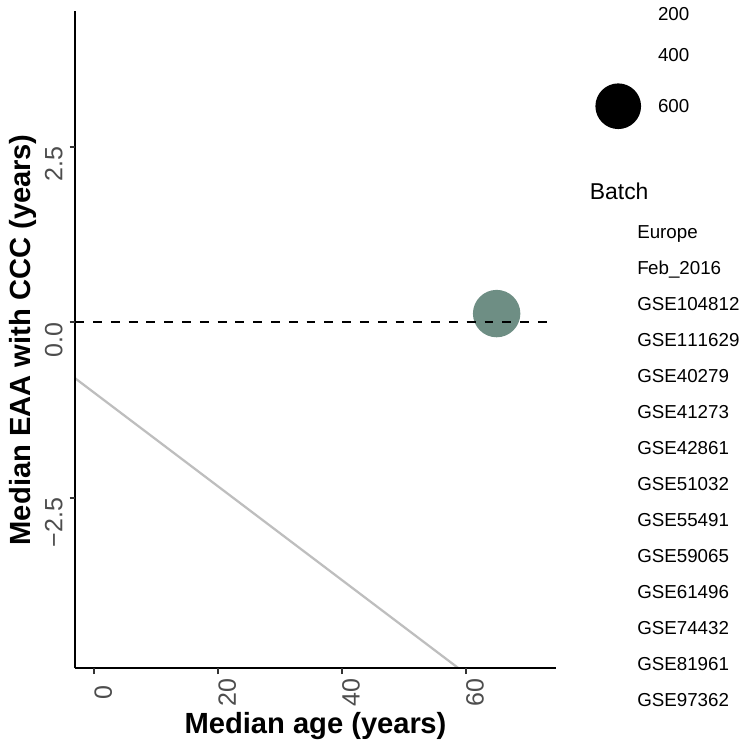

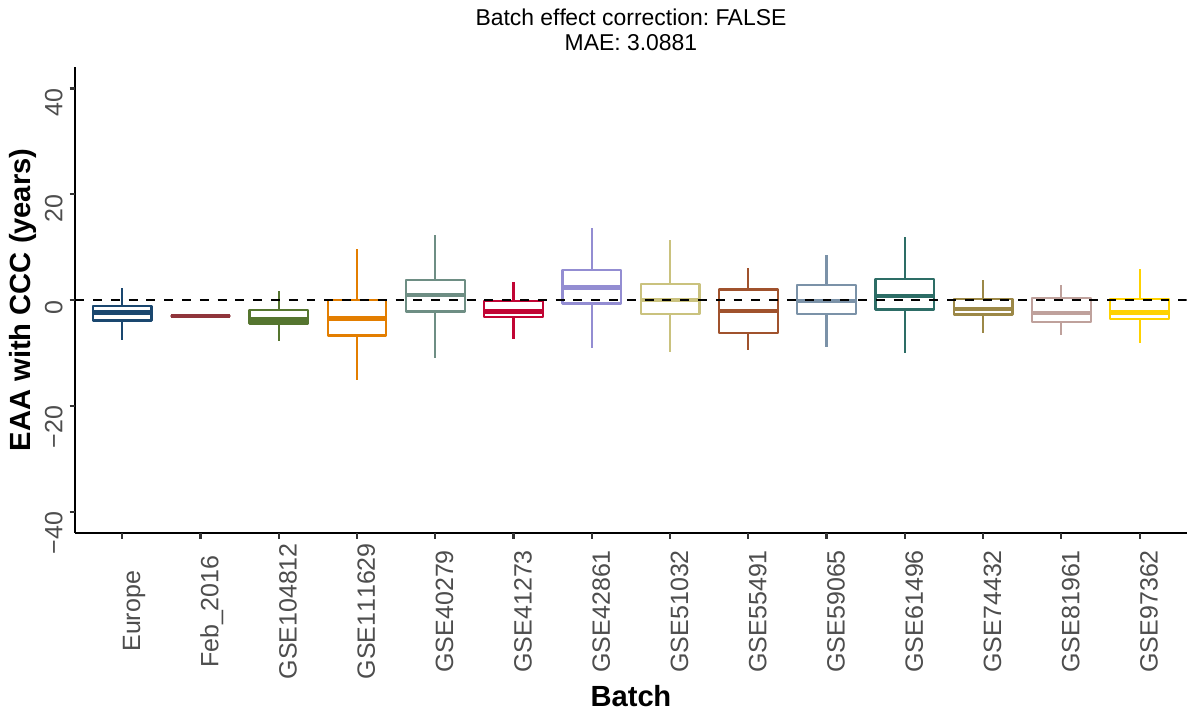

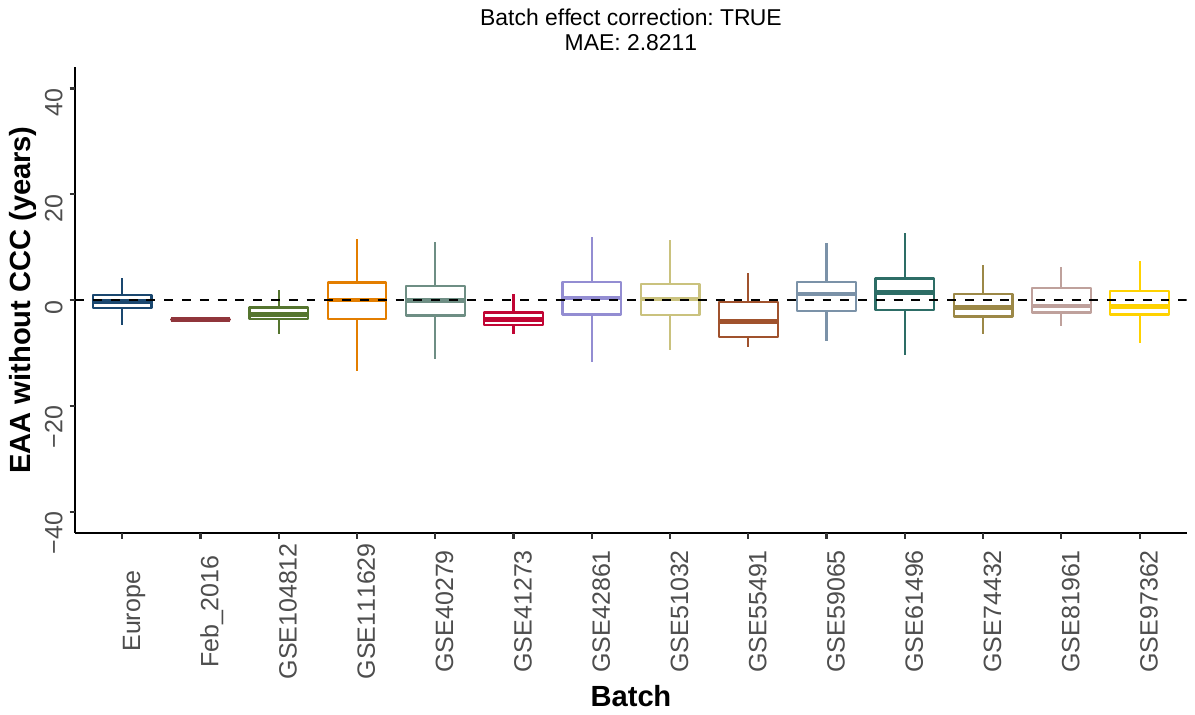

**d**

**Figure S1.** Complementary to Fig. 1. **a.** Distribution of the epigenetic age acceleration (EAA) without cell composition correction (CCC) for the different control batches, before applying batch effect correction. MAE: median absolute error. **b.** Distribution of the EAA with cell composition correction (CCC) for the different control batches, before applying batch effect correction. **c.** Scatterplot showing the values of the first two principal components (PCs) for the cases (developmental disorder samples) after performing PCA on the control probes of the 450K arrays. Each point corresponds to a different case sample and the colours represent the different batches. The different batches cluster together in the PCA space, showing that the control probes indeed capture technical variation. Please note that all the PCA calculations were done with more samples from cases and controls than those that were included in the final screening since it was performed before the filtering step (see Methods). **d.** Plot showing the percentages of technical variance explained by the different PCs from the control probes. The dashed line represents the optimal number of PCs (17) that was finally used. **e.** Distribution of the EAA without cell composition correction (CCC) for the different control batches, after applying batch effect correction. **f.** After batch effect correction, deviations from a median EAA of zero (dotted black line) in some of the control batches can be explained by other causes. The grey line separates in the lower left corner those weird batches (Feb_2016, GSE104812, GSE41273, GSE55491), which have a small sample size and/or a low median age.

**Figure S2**

**a**

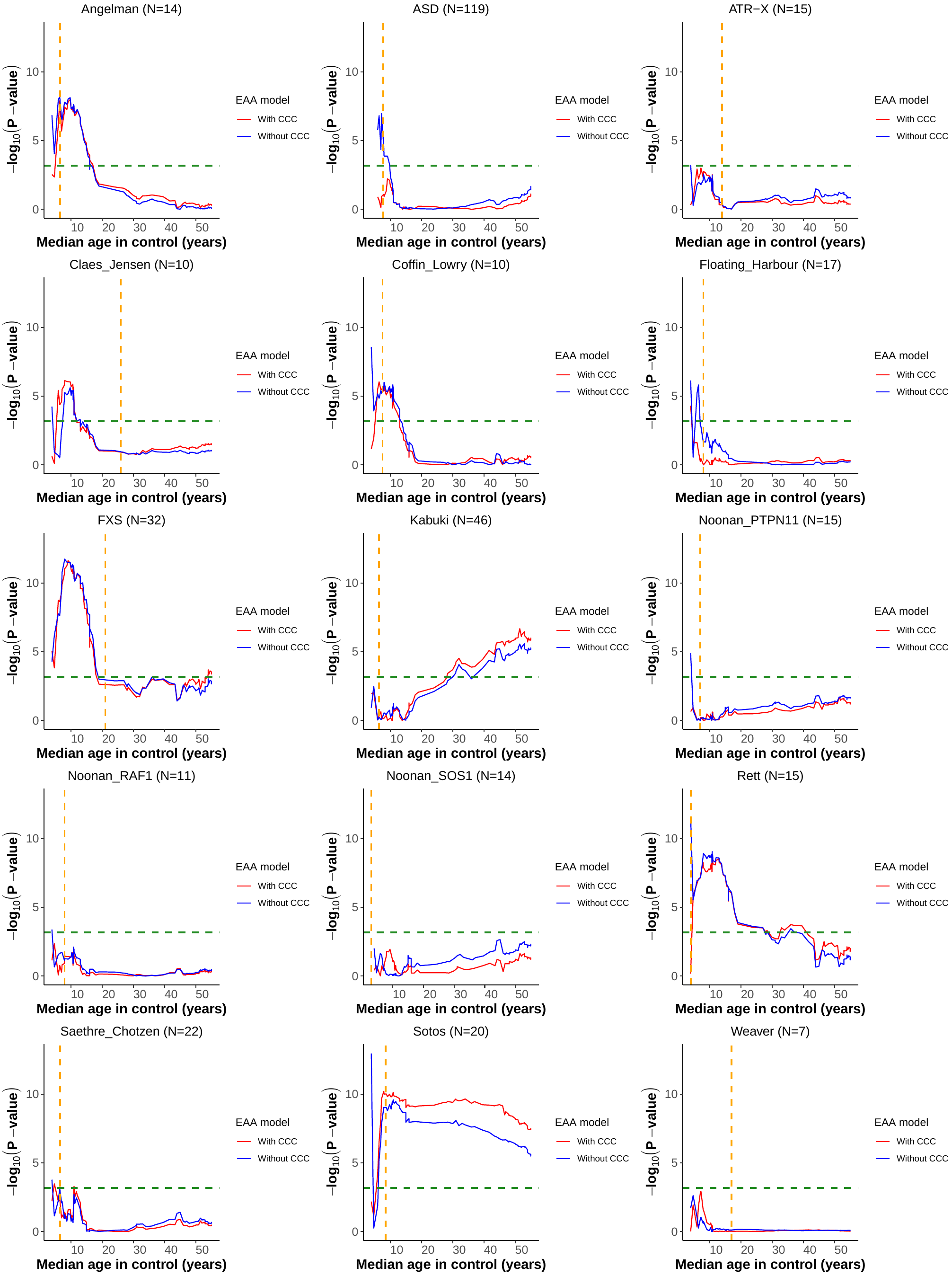

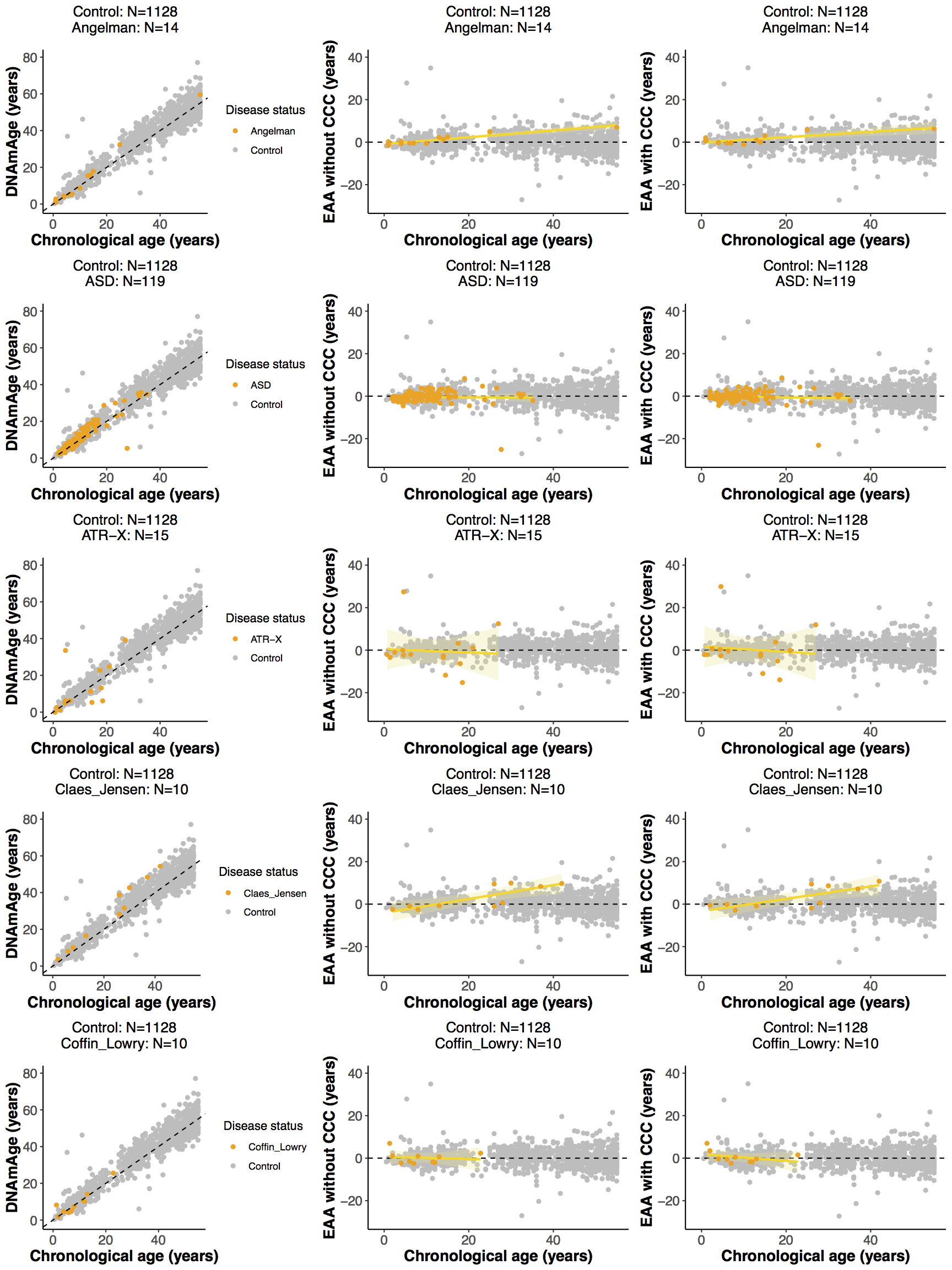

**b**

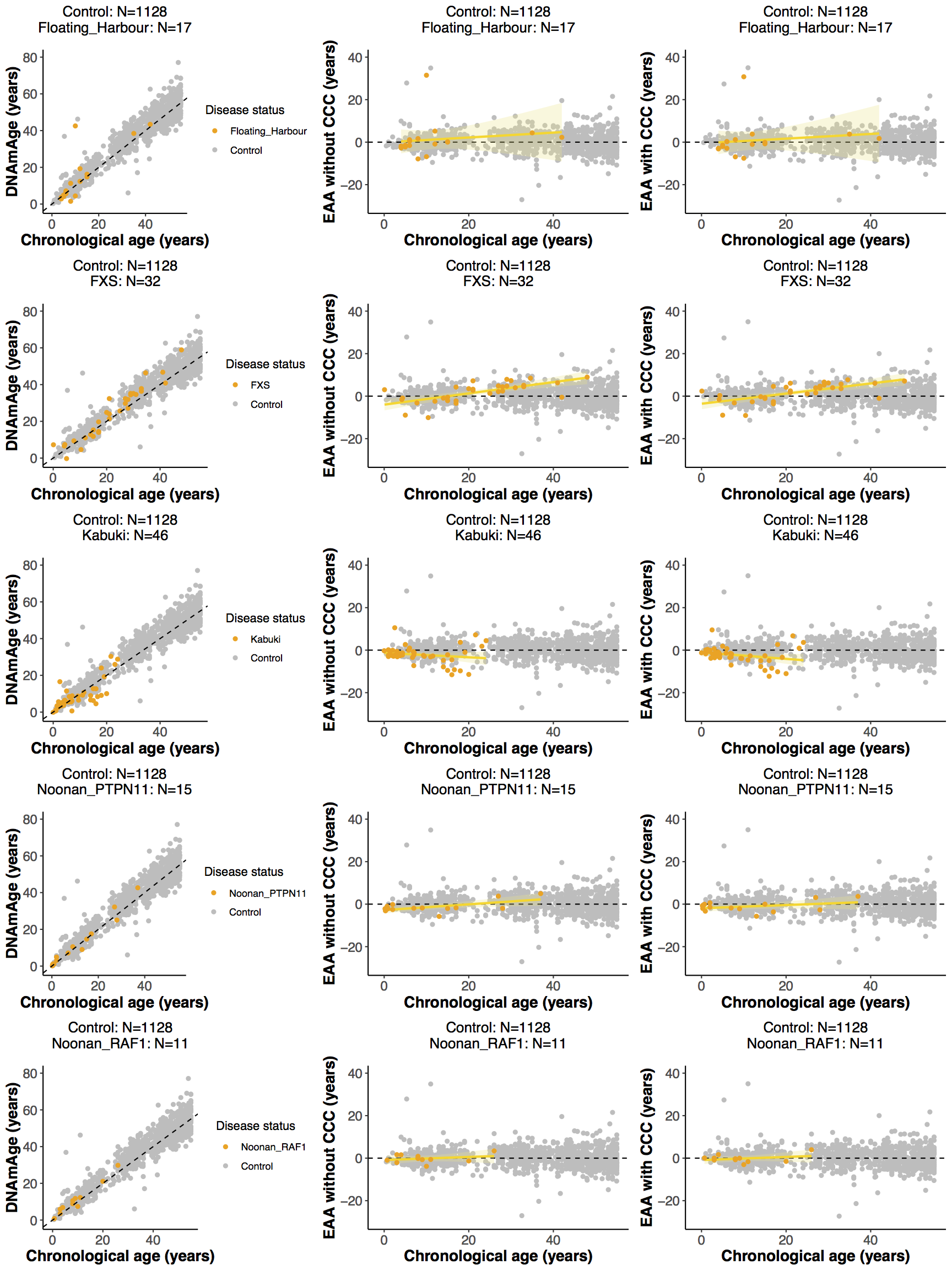

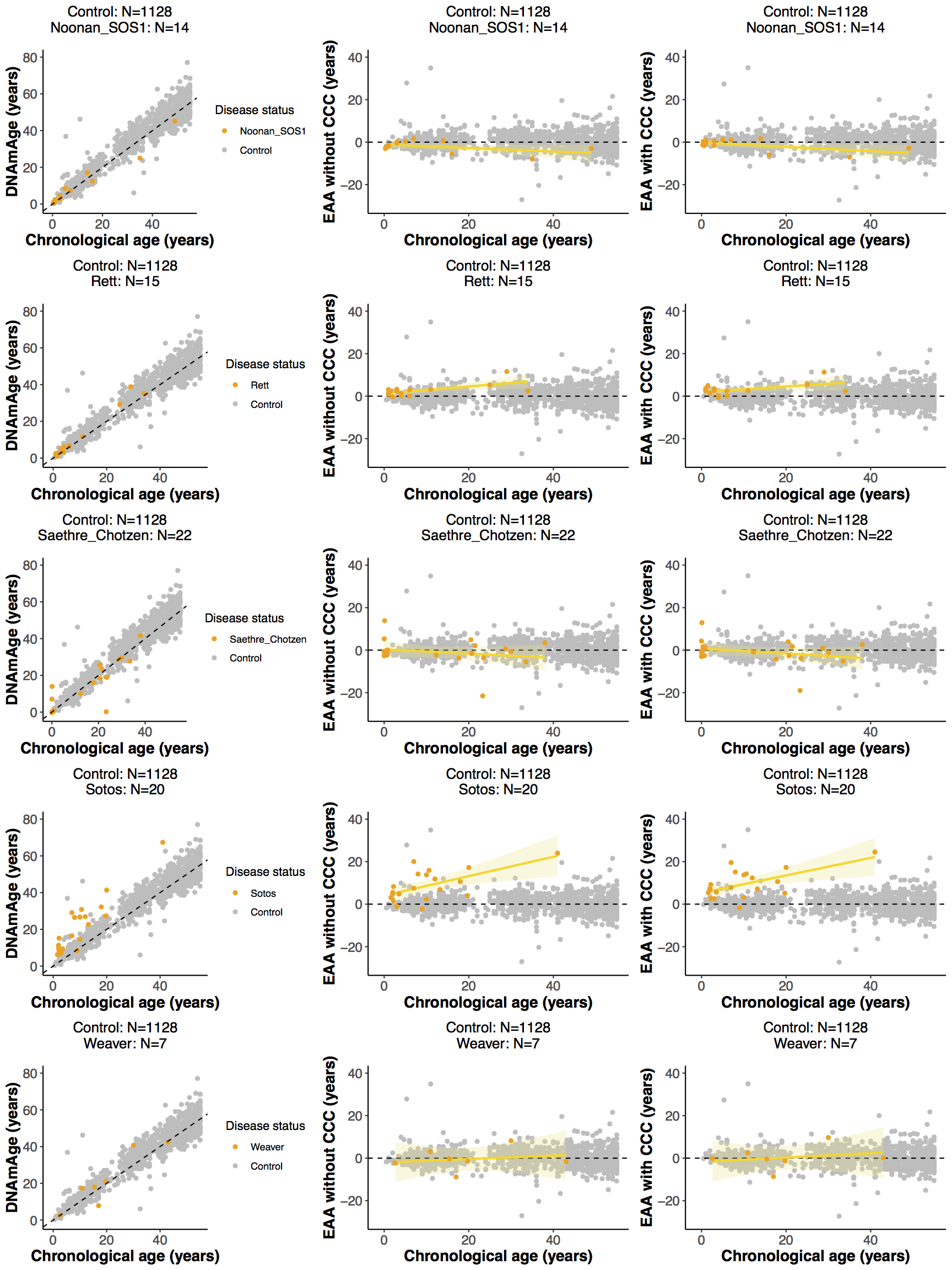

**Figure S2.** Complementary to Fig. 2. **a.** Effect of changing the median age of the controls when performing the screening for epigenetic age acceleration (EAA) in the different developmental disorders. The dashed green line displays the significance level of α = 0.01 after Bonferroni correction. The dashed orange line displays the median age for the samples in the developmental disorder considered. In blue: EAA model without cell composition correction (CCC). In red: EAA model with CCC. **b.** Left panel: scatterplot showing the relation between epigenetic age (*DNAmAge*) according to Horvath’s model and chronological age of the samples for a given developmental disorder (orange) and control (grey). Each sample is represented by one point. The black dashed line represents the diagonal to aid visualisation. Middle and right panels: scatterplots showing the relation between the epigenetic age acceleration (EAA) (without and with CCC respectively) and chronological age of the samples for a given developmental disorder (orange) and control (grey). Each sample is represented by one point. The yellow line represents the linear model *EAA ~ Age*, with the standard error shown in the light yellow shade.

**Figure S3**

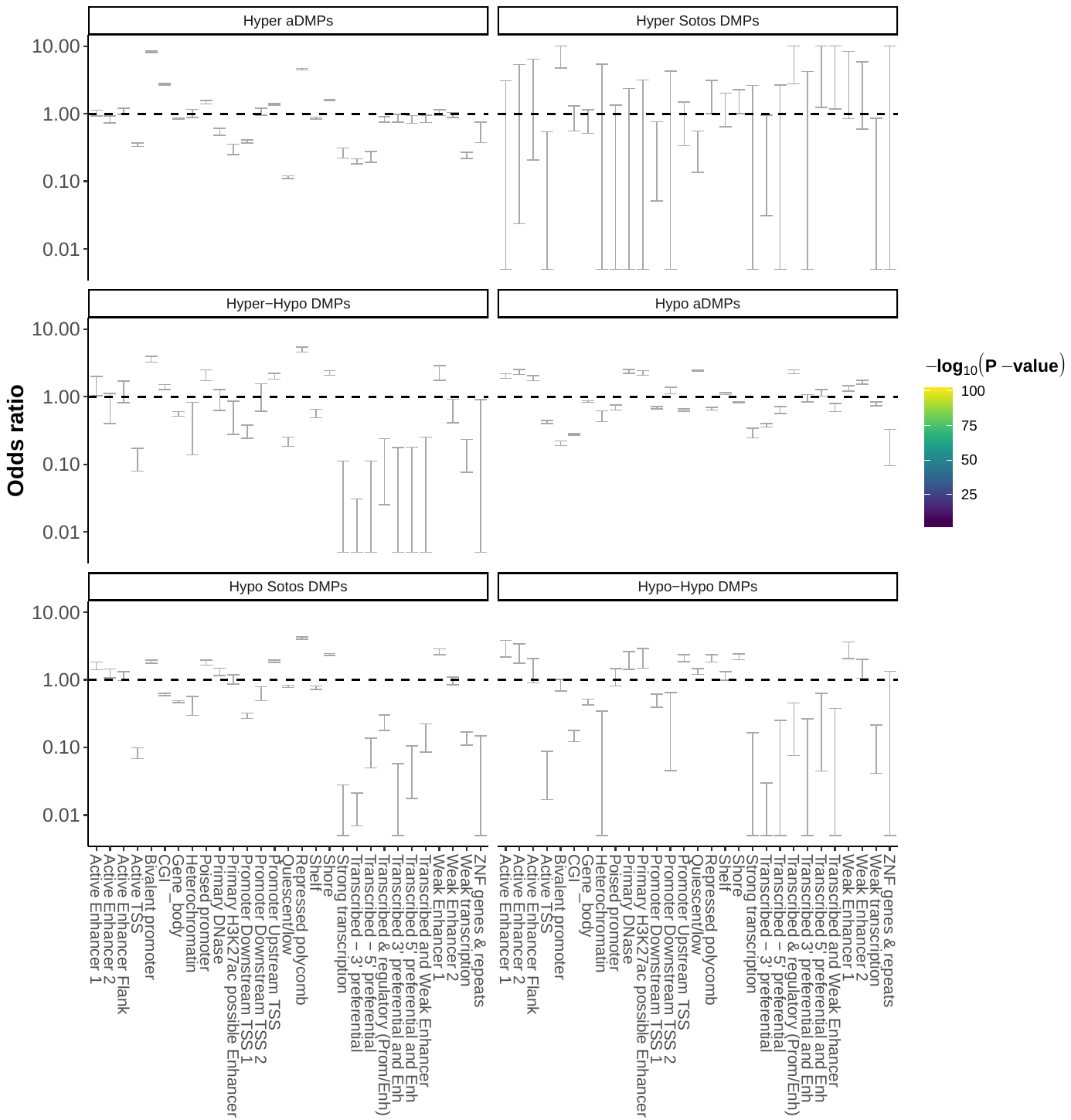

**a**

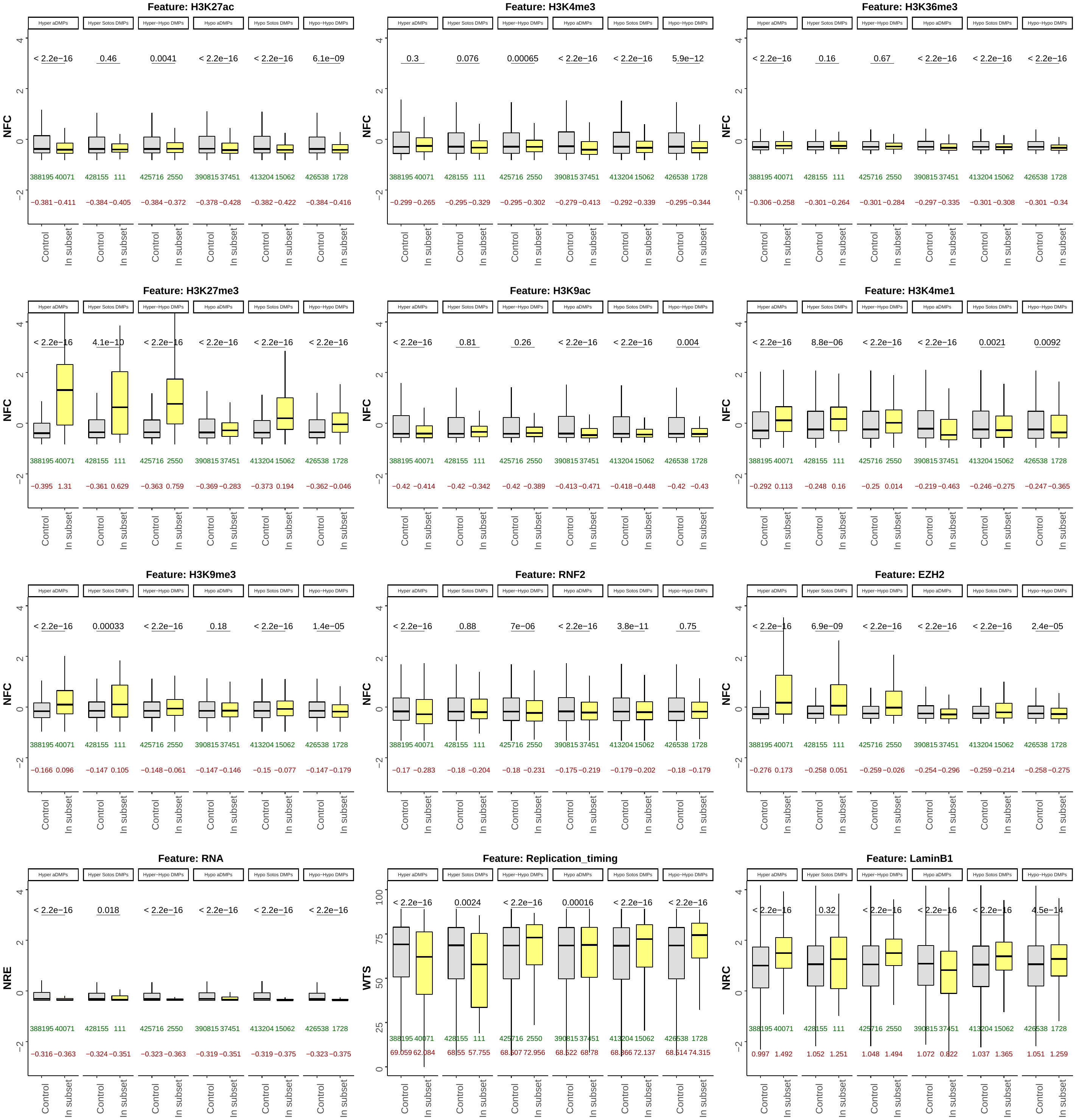

**b**

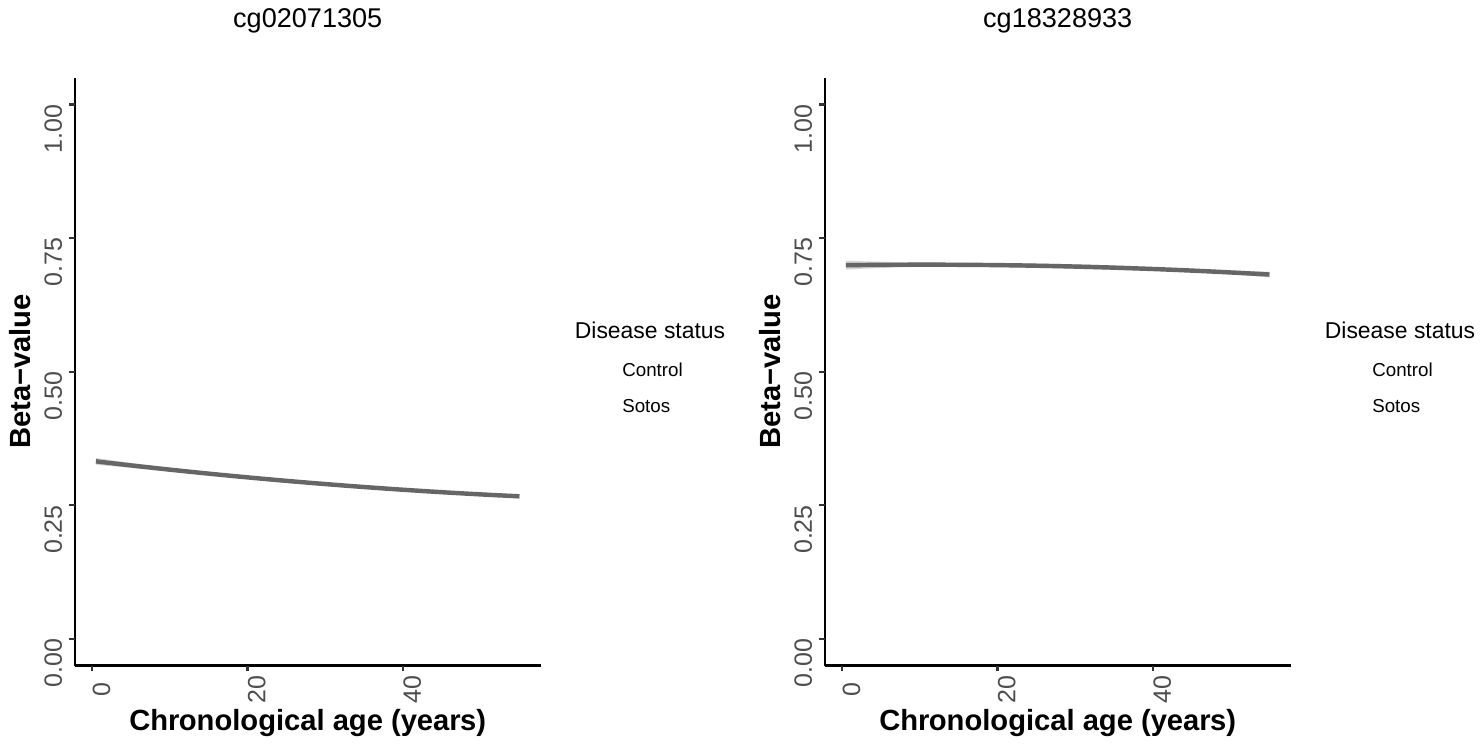

**c**

**d**

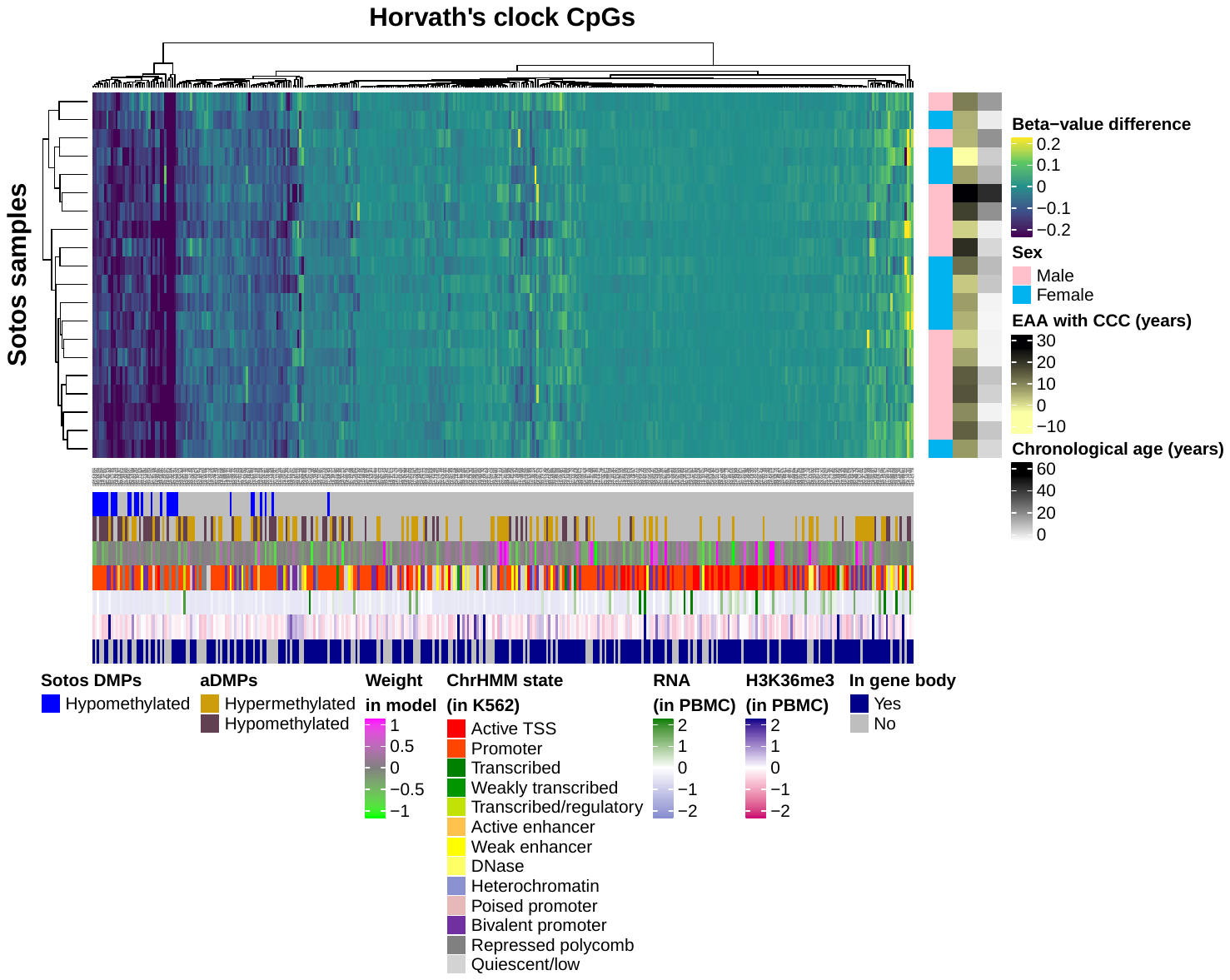

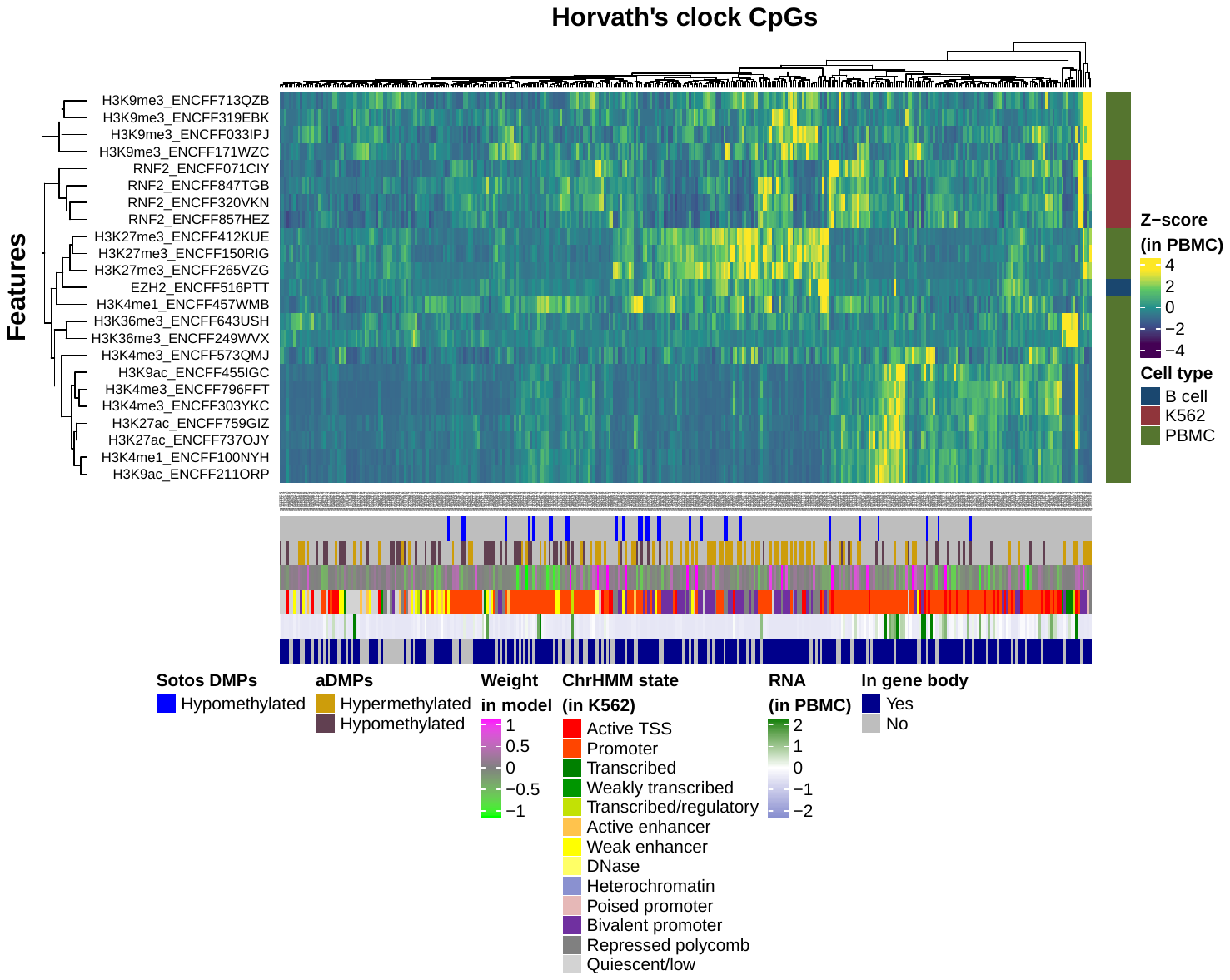

**e**

**f**

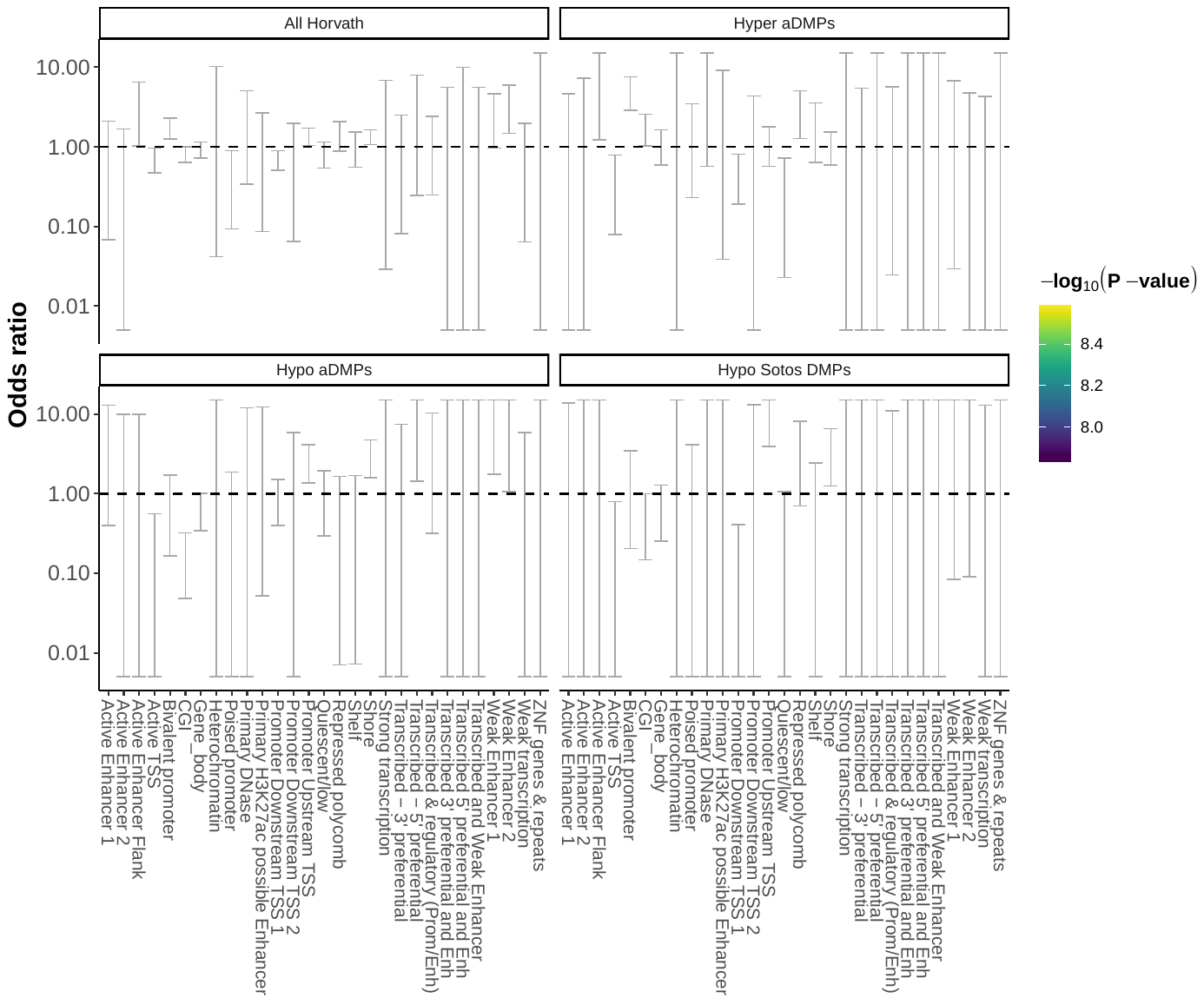

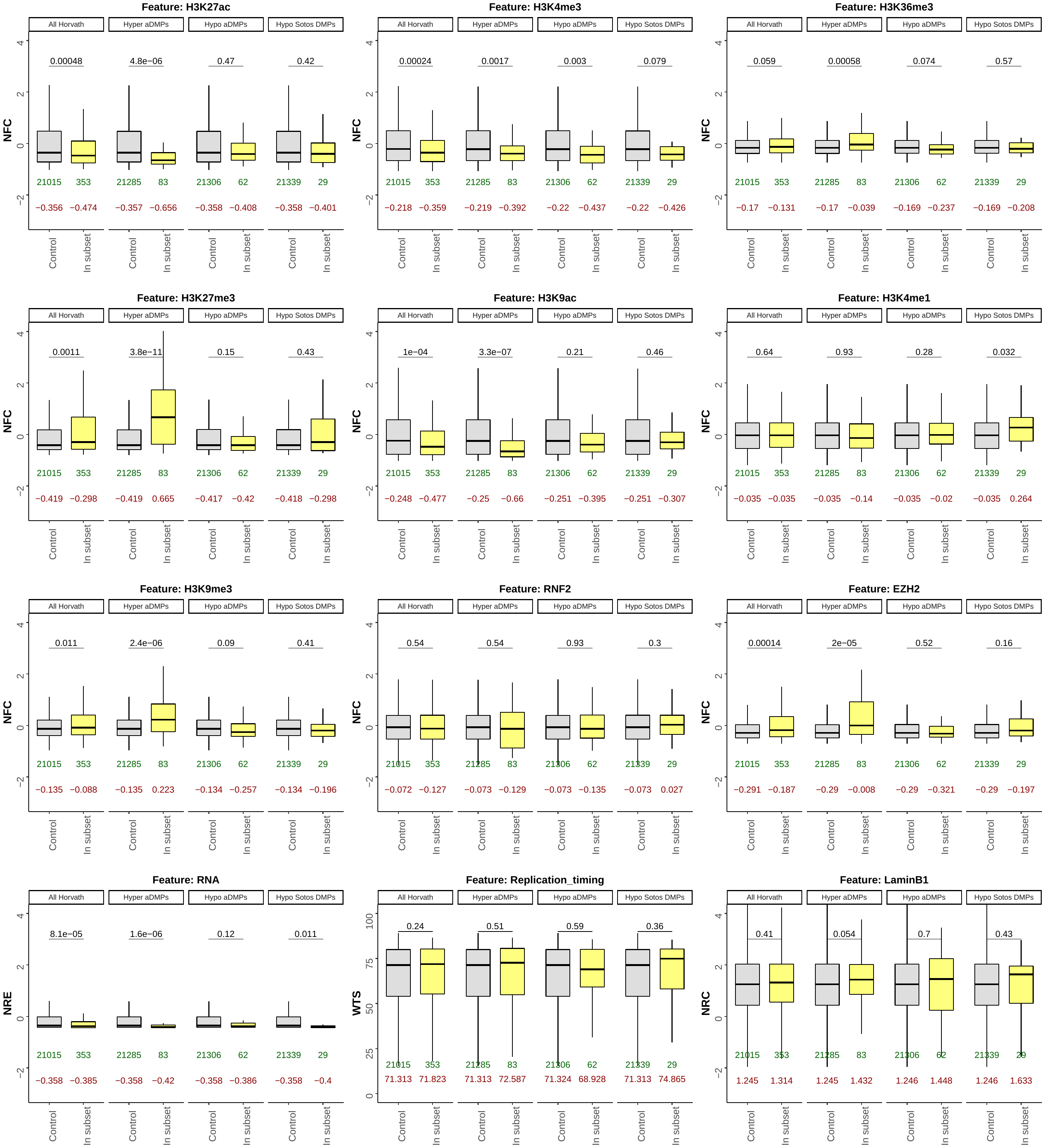

**g**

**Figure S3.** Complementary to Fig. 3. **a.** Enrichment for the categorical (epi)genomic features considered when comparing the different genome-wide subsets of differentially methylated positions (DMPs) in ageing and Sotos against a control (see Methods). The y-axis represents the odds ratio (OR), the error bars show the 95% confidence interval for the OR estimate and the colour of the points codes for -log_10_(p-value) obtained after testing for enrichment using Fisher’s exact test. An OR > 1 shows that the given feature is enriched in the subset of DMPs considered, whilst an OR < 1 shows that it is found less than expected. The ‘Hyper-Hypo DMPs’ subset results from the intersection between the hypermethylated DMPs in ageing and the hypomethylated DMPs in Sotos. The ‘Hypo-Hypo DMPs’ subset results from the intersection between the hypomethylated DMPs in ageing and Sotos. In grey: features that did not reach significance using a significance level of α = 0.01 after Bonferroni correction. **b.** Boxplots showing the distributions of scores (see Methods) for the continuous (epi)genomic features considered when comparing the different genome-wide subsets of differentially methylated positions (DMPs) in ageing and Sotos against a control (see Methods). The p-values (two-sided Wilcoxon’s test, before multiple testing correction) are shown above the boxplots. The number of DMPs belonging to each subset (in green) and the median value of the feature score (in dark red) are shown below the boxplots. NFC: ‘normalised fold change’; NRE: ‘normalised RNA expression’; WTS: ‘wavelet-transformed signals’; NRC: ‘normalised read counts’. **c.** DNA methylation (beta-value) profiles for two of the clock CpG sites (cg02071305 and cg18328933). A linear model (displayed in dark grey) can be fixed to each CpG site to model the changes in beta-value with chronological age in the controls (grey). Afterwards, the difference of the Sotos samples beta-values (orange) with the controls can be estimated. **d.** Heatmap displaying the differential methylation patterns for Sotos samples (rows) when compared with controls in each one of the 353 clock CpGs (columns). Hierarchical clustering was performed in both rows and columns. RNA refers to the ‘normalised RNA expression’ (NRE, see Methods). H3K36me3 refers to the H3K36me3 histone modification ‘normalised fold change’ (NFC, see Methods). aDMPs: differentially methylated positions during ageing. EAA: epigenetic age acceleration. CCC: cell composition correction. PBMC: peripheral blood mononuclear cells. **e.** Heatmap displaying the scores for the different continuous (epi)genomic features (rows) in each one of the 353 clock CpGs (columns). The names of the features include the ENCODE ID. Hierarchical clustering was performed in both rows and columns. **f.** Same as a., but focused on the 353 clock CpG sites. **g.** Same as b., but focused on the 353 clock CpG sites.

**Figure S4**

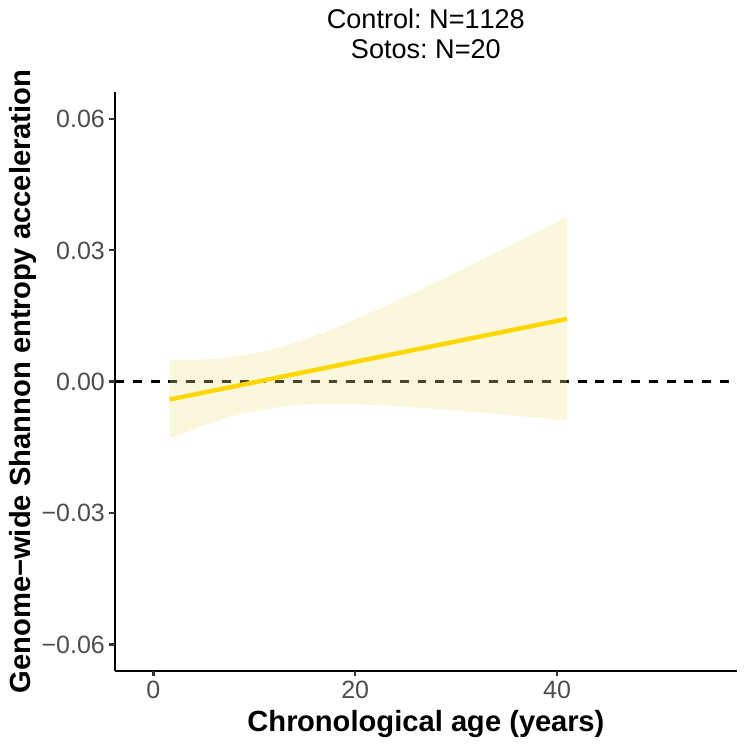

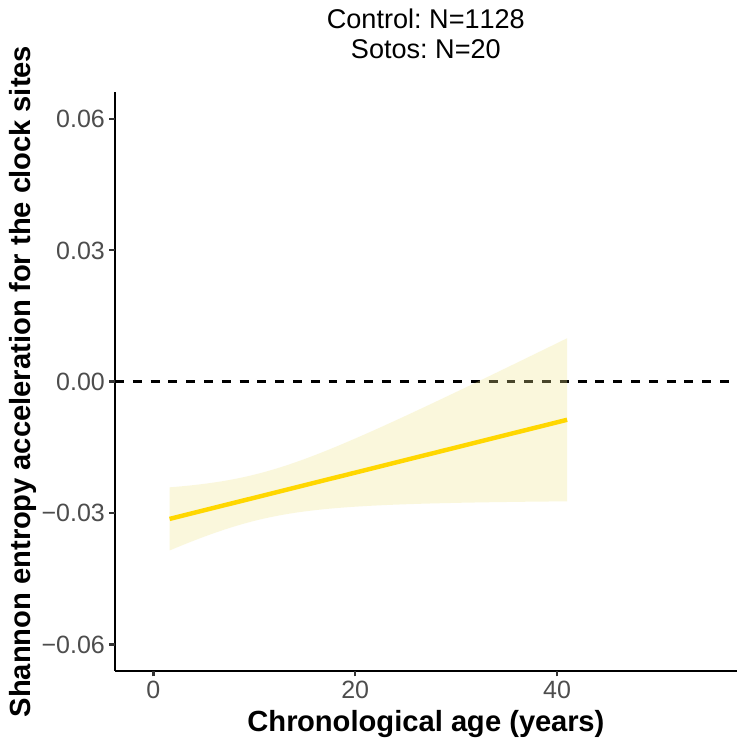

**a**

**b**

**c**

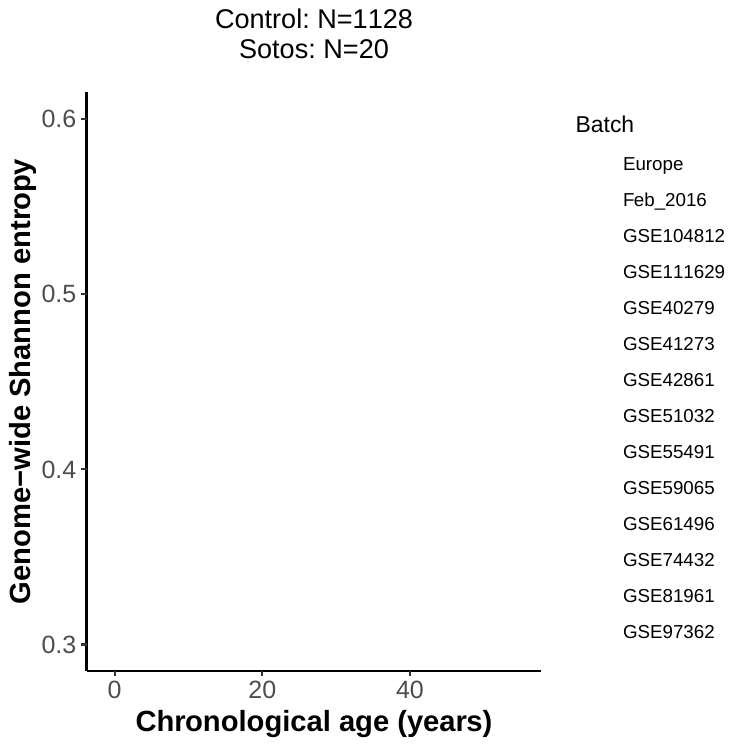

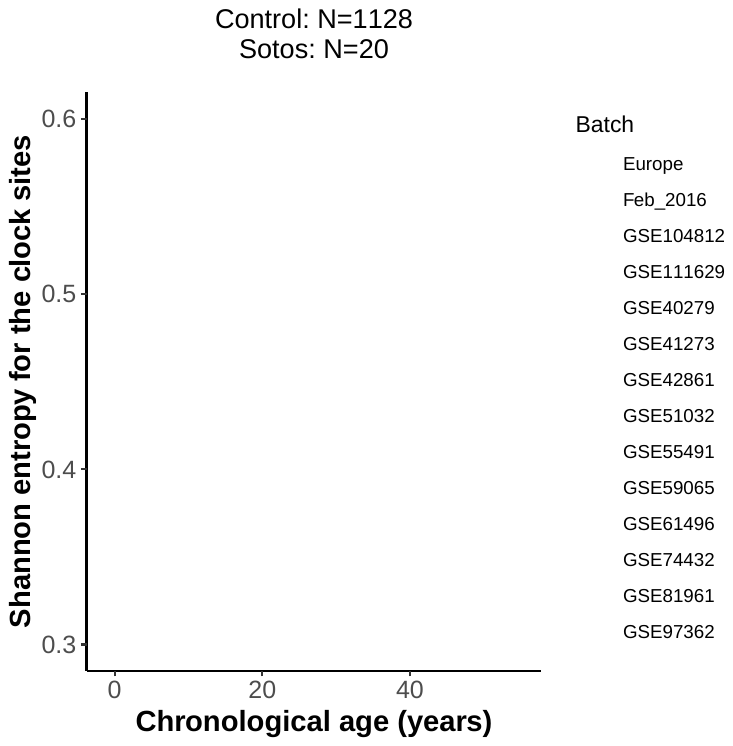

**d**

**Figure S4.** Complementary to Fig. 4. **a.** Scatterplot showing the relation between the genome-wide Shannon entropy acceleration (*gSEA*) and chronological age of the samples for Sotos (orange) and healthy controls (grey). Each sample is represented by one point. The yellow line represents the linear model *gSEA ~ Age*, with the standard error shown in the light yellow shade. **b.** Same as a., but using the Shannon entropy acceleration calculated only for the 353 CpG sites in the Horvath epigenetic clock (*cSEA*). **c.** Scatterplot showing the effects of the different batches on the genome-wide Shannon entropy calculations. Each sample is represented by one point and coloured according to the batch that they belong to. **d.** Same as c., but using the Shannon entropy calculated only for the 353 CpG sites in the Horvath epigenetic clock.

**Figure S5**

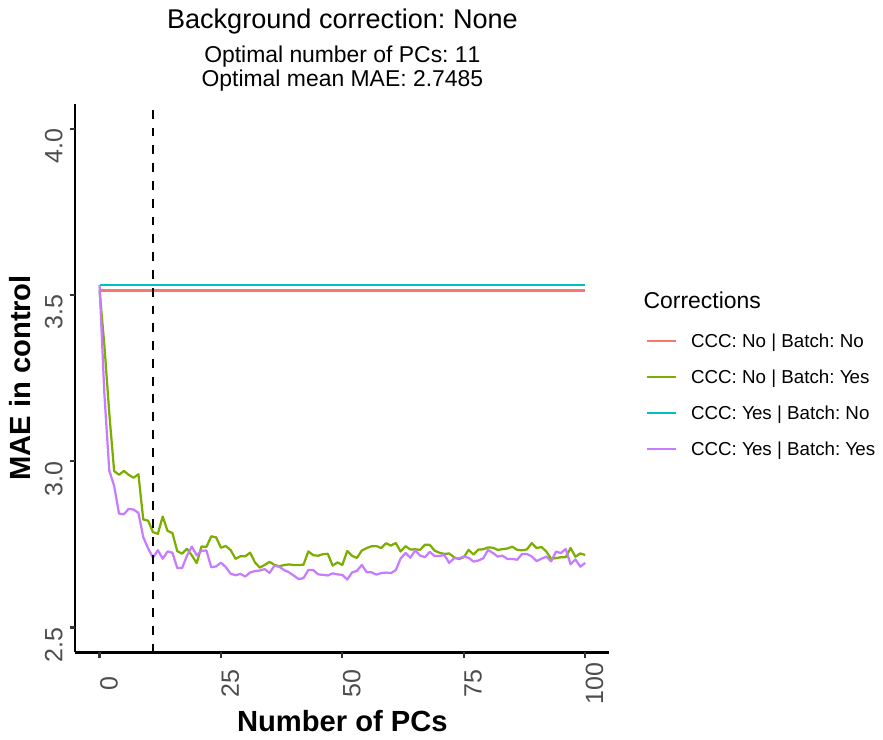

**b**

**a**

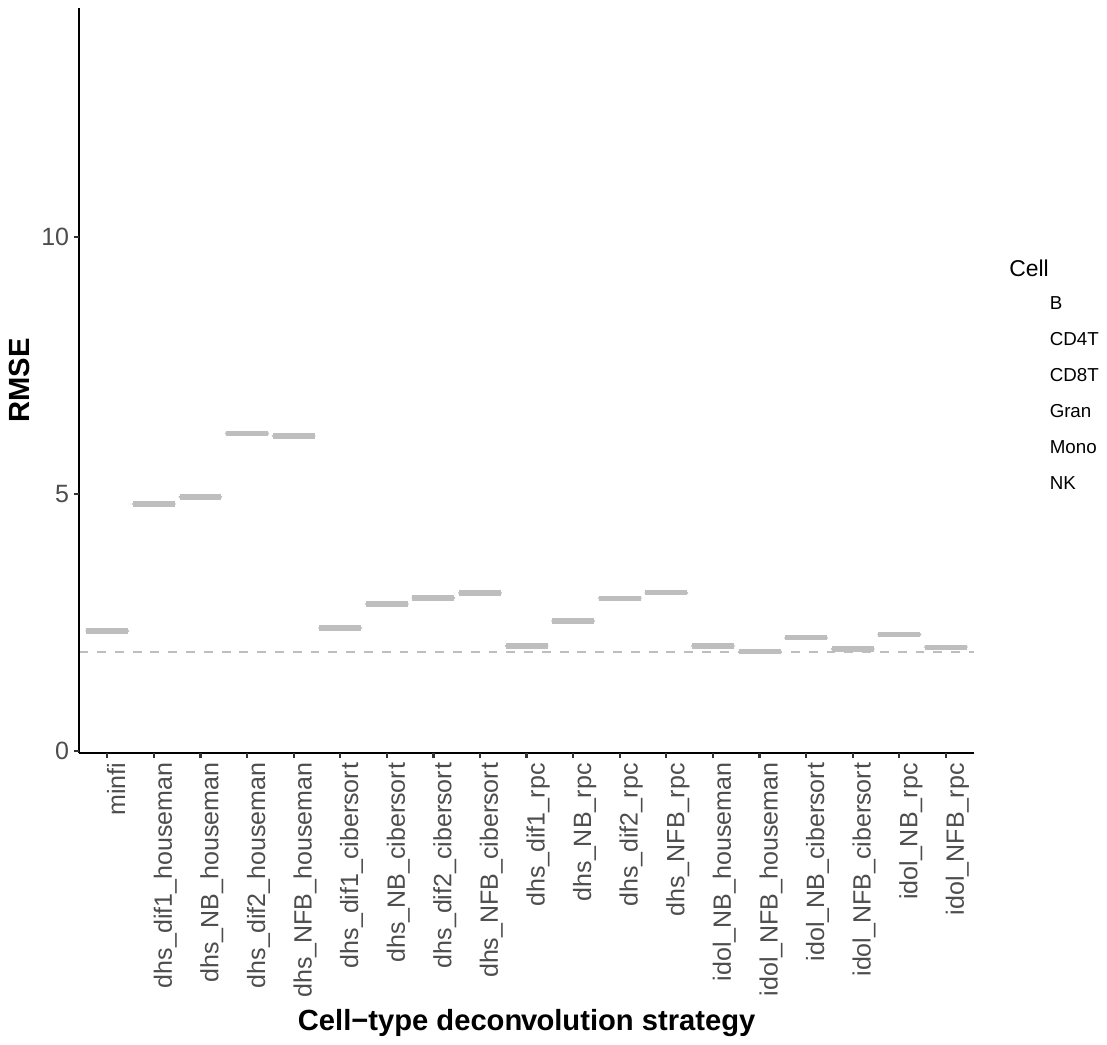

**Figure S5.** Complementary to Methods. **a.** Plot showing how the median absolute error (MAE) of the prediction in the control samples, that should tend to zero, is reduced when the PCs capturing the technical variation are included as part of the modelling strategy (see Methods). In this case, no background correction was performed, as opposed to the results in Fig. 1c. The dashed line represents the optimal number of PCs (17) that was finally used. The optimal mean MAE is calculated as the average MAE between the green and purple lines. CCC: cell composition correction. **b.** Benchmarking of the different strategies for cell-type deconvolution in blood. The root mean square error (RMSE) quantifies the deviations between our estimates and the actual proportions of cells for a gold-standard dataset (GSE77797). The description for the different strategies can be found in Additional file 4.
