## Supplementary material for "Screening for genes that accelerate the epigenetic ageing clock in humans reveals a role for the H3K36 methyltransferase NSD1": additional_files_legends.docx

**Additional file 1.** Supplementary figures that complement the main manuscript.

**Additional file 2.** Information for the samples with developmental disorders (cases) that were included in the main screen (N=367).

**Additional file 3.** Information for the healthy control samples that were included in the main screen (N=1128).

**Additional file 4.** Information about the different blood cell-type deconvolution strategies that were benchmarked against the gold-standard dataset (GSE77797).

**Additional file 5.** Information (including the source) about the continuous (epi)genomic features (ChIP-seq and RNA-seq data) that were included in our analysis to annotate the different sets of CpG sites.
